# HIV-1 Capsid Shape, Orientation, and Entropic Elasticity Regulate Translocation into the Nuclear Pore Complex

**DOI:** 10.1101/2023.08.05.552137

**Authors:** Arpa Hudait, Gregory A. Voth

## Abstract

Nuclear import of the viral capsid is a critical step in the HIV-1 life cycle that serve to transport and release genomic material into the nucleus. Nuclear Pore Complex (NPC) allows passage of intact capsid, though mechanistic details of the process remain to be fully understood. Here we investigate the factors regulating HIV-1 capsid translocation into the NPC central channel using coarse-grained molecular dynamics simulations. We find that successful translocation is contingent on the compatibility of the capsid morphology and channel dimension and the proper orientation of the capsid approach to the channel. The central channel dynamically expands to allow capsid passage, demonstrating the pleomorphic nature of the channel necessary for transporting large cargoes. Structural analysis shows that stress induced by the central channel confinement and uncondensed internal genomic material generates correlated striated patterns of lattice disorder across the viral capsid surface which is an indicator of its lattice “elasticity”. Our results suggest that the “elasticity” can aid the capsid to adapt to the stress and remain structurally intact during translocation.

**Teaser:** Computer Simulations identify capsid elasticity as a key factor for successful nuclear entry.

## INTRODUCTION

HIV-1 spreads infection through a replication cycle that comprises several distinct steps. The transport of the HIV-1 genome into the nucleus is an imperative step in the life cycle for viral infection (1). The HIV-1 mature capsid encloses and protects the genomic RNA and the critical enzymes for replication, reverse transcriptase, and integrase (2, 3). The HIV-1 mature capsid comprises 1000-1500 copies of the capsid (CA) proteins forming a predominantly hexameric lattice while incorporating 12 pentamers during its assembly to achieve closure (4-7). The typical capsid cone is of length ∼120 nm and width ∼60 nm (8, 9). Nuclear import of HIV-1 capsid is mediated through the nuclear pore complex (NPC) embedded in the nuclear envelope. Early structural studies determined the diameter of the NPC central channel that mediates cargo transport to be ∼ 40 nm (10). Due to the incompatibility in size between the HIV-1 capsid cone and NPC central channel, initial models of viral nuclear import and capsid uncoating proposed that uncoating of the capsid is initiated in the cytoplasm or when the capsid is docked at the NPC (11-15). In contrast to the early models, recent experiments tracking the HIV-1 cores in live infected cells demonstrated that intact capsids are transported through the nuclear pore, reverse transcription occurs in the (at least largely) intact capsid, and uncoating occurs before integration in the vicinity of the genomic integration site (16, 17). This observation is also supported by direct visualization of capsids through cryo-electron tomography (cryo-ET) and electron microscopy techniques that revealed the transport of intact capsids across the nuclear pore (18, 19). At present, however, it is not known what fraction of the total capsids present in an infected cell can translocate into and through the nuclear pore, nor what structural and other characteristics the successful ones may have. Moreover, these recent findings also suggest that nuclear pore complex (NPC) can exist in a dilated state allowing translocation of intact HIV-1 capsids. How the physical properties of capsid lattice are impacted during passage into the NPC central channel is also yet to be fully understood and can provide valuable insight into the molecular mechanism of capsid structural failure and rupture after entry to the nucleus.

The NPC, with a molecular weight of ∼120 MDa and consisting of up to 1000 proteins, is one of the largest protein assemblies in the cell. Several recent studies have determined the structure of human NPC in unprecedented detail, allowing for a detailed view of previously unassigned segments of the macromolecular complex (20-22). The constituent proteins of the NPC are called nucleoporins (NUPs). NUPs assemble to form hetero-oligomeric complexes establishing three stacked concentric rings in an eightfold rotational symmetry. Therefore, each ring consists of eight spokes. The building block of the outer cytoplasmic ring (CR) and nuclear ring (NR) is the Y-complex oligomerized in a head-to-tail arrangement forming two concentric and slightly shifted eight-membered rings (10). Similarly, multiple copies of NUPs comprise a single spoke of the inner ring (IR) complex. Importantly, linker NUPs interconnect adjacent spokes of the IR complex allowing flexibility in the relative arrangement of spokes. Key for the nucleocytoplasmic exchange of cargo is the intrinsically disordered phenylalanine-glycine (FG) motifs tethered to the NPC scaffold. The HIV-1 capsid is known to interact with several FG-containing NUPs at the CR, IR, and NR (23-25).

The NPC is one of the largest macromolecular complexes in the cell. Simulations of membrane-embedded NPC with viral capsid at atomistic detail would require over a billion atoms and is presently and in the foreseeable future computationally infeasible due to the spatial and temporal scale of the multistep translocation process. Using systematically derived “bottom-up” CG models, one can instead perform physically reliable coarse-grained (CG) molecular dynamics (MD) simulations of large protein complexes at scales of relevance to typical cellular processes. Here, “bottom-up” signifies that CG molecular model and interactions are systematically constructed from underlying atomistic interactions (26). In other words, the derived CG model and interactions are designed to reproduce the molecular behavior sampled in the atomistic simulations when the latter are projected exactly onto a coarser representation. As leading example, CG MD simulations in our group’s hands have been particularly effective in uncovering essential mechanistic details of large-scale viral processes, such as factors that regulate HIV-1 capsid lattice growth and formation (7, 27-29), capsid restriction (30), and immature Gag assembly (31-33), as well as the concerted interactions that guide the SARS-CoV-2 virus for membrane fusion and entry (34). Hence in this work, we first derive a “bottom-up” CG molecular model and interactions of NUP monomer and subcomplexes from the recent high-resolution structural data (PDB: 7R5K and 7R5J) of human NPC (20). Additionally, we also develop a CG molecular model of HIV-1 CA monomer (the same resolution as NPC) and the CG associative interactions between CA-CA and CA-FG. The CG model of NUP monomers and subcomplexes are combined to generate a composite model of constricted and dilated membrane-embedded human NPC. The composite CG NPC model developed in this work consists of the outer CR and NR, IR, and the NUPs containing the capsid-binding FG-repeats at the IR (NUP54, NUP58, and NUP62). We created another composite NPC model in which the disordered NUP98 chains are also tethered to the IR. The CG models in this work are the so-called “solvent free” CG models in which the effects of the solvent are folded into the CG interactions. Similar models are sometimes referred to as “implicit solvent” models, but that phrase can carry a specific connotation as to how the effects of solvent are included and often such models are expressed at full atomic resolution.

In this work, we elucidate the HIV-1 capsid translocation dynamics into the NPC central channel using CG MD simulations. Specifically, we simulate the translocation dynamics of three different capsid morphologies into the central channel of constricted and dilated states of NPC using the composite CG NPC and HIV-1 capsid CG models. Our simulations show that the dilated state of NPC allows translocation of the cone-shaped capsid into the NPC when docked at the narrow end. The dilated state of NPC also allows translocation of a pill-shaped capsid. The constricted NPC impedes the translocation of all capsid morphologies examined in the simulations. We find that capsid translocation is driven by the energetics of the interaction between the capsid binding central channel FG-NUPs. In contrast, incompatibility between the channel size and capsid morphology is the primary barrier to nuclear entry. Analysis of the viral capsid structures docked at the NPC further reveals the appearance of distinct striated patterns of lattice disorder along the surface of the capsid. This lattice disorder at the same time represents a certain fragility of a more perfect capsid lattice but also a form of capsid “flexibility” in terms of an entropic “spring-like” response to the stress imposed on the capsid by the NPC. To mimic the initiation of reverse transcription, we also introduce a model for the viral genomic complex inside the capsids docked at the NPC. We find that in the uncondensed form, the genomic complex significantly amplifies the structural fragility of the pill-shaped capsid compared to the more canonical cone-shaped capsid.

Overall, our analysis of the intact viral capsids docked at the NPC demonstrates that the capsid lattice is pliable, but the structural integrity is also weakened during passage through the NPC central channel. The findings reported in this work elucidate several key factors that regulate the successful passage of viral capsid from the cytoplasmic to the nuclear end of the NPC.

## RESULTS

### Coarse-grained model of composite membrane-embedded human NPC

We developed CG models of the constricted and dilated states of human NPC from available experimental structural and biochemical data to surmount the computational costs associated with simulating a large macromolecular complex in atomic detail (20). The composite CG NPC models presented in this work consist of the outer CR and NR, and IR (**Fig. 1*A*** and **1*B***). As noted earlier, the model development approach is largely “bottom-up”, i.e., constructed quantitatively from the underlying atomistic interactions generated from extensive all-atom (AA) molecular dynamics (MD) simulation trajectories of NUP monomers and heterodimer subcomplexes. The statistics from the AA MD simulation trajectories of NUP monomers were used to map CG molecular models with a CG mapping resolution of ∼ 5 amino acids per CG site or “bead” (35). The same AA MD trajectories were used to derive a heterogeneous elastic network model (hENM) of effective harmonic interactions to represent the intra-monomer interactions and maintain the protein molecular shape (36). Additionally, each CG bead had an excluded volume that allowed for maintaining the protein shape and preventing unphysical overlap between neighboring beads. Inter-monomer interactions within a subcomplex were phenomenologically modeled with a bonded soft elastic network model (harmonic force constant of 0.01 kcal mol^−1^ Å^−2^) derived directly from the cryo-ET structure. The choice of a weak force constant allows significant configurational flexibility within a subcomplex while maintaining the overall shape of the subcomplexes. Finally, inter-protein associative interactions at key binding interfaces between different NUP subcomplexes were modeled using non-bonded short-ranged attractive interactions derived from AA MD simulations of the corresponding heterodimer NUP complexes (37). Details of all the AA MD simulations, model parameterization procedures, and complete CG model details are provided in the *Methods* and *Supporting Information*.

**Fig. 1.**
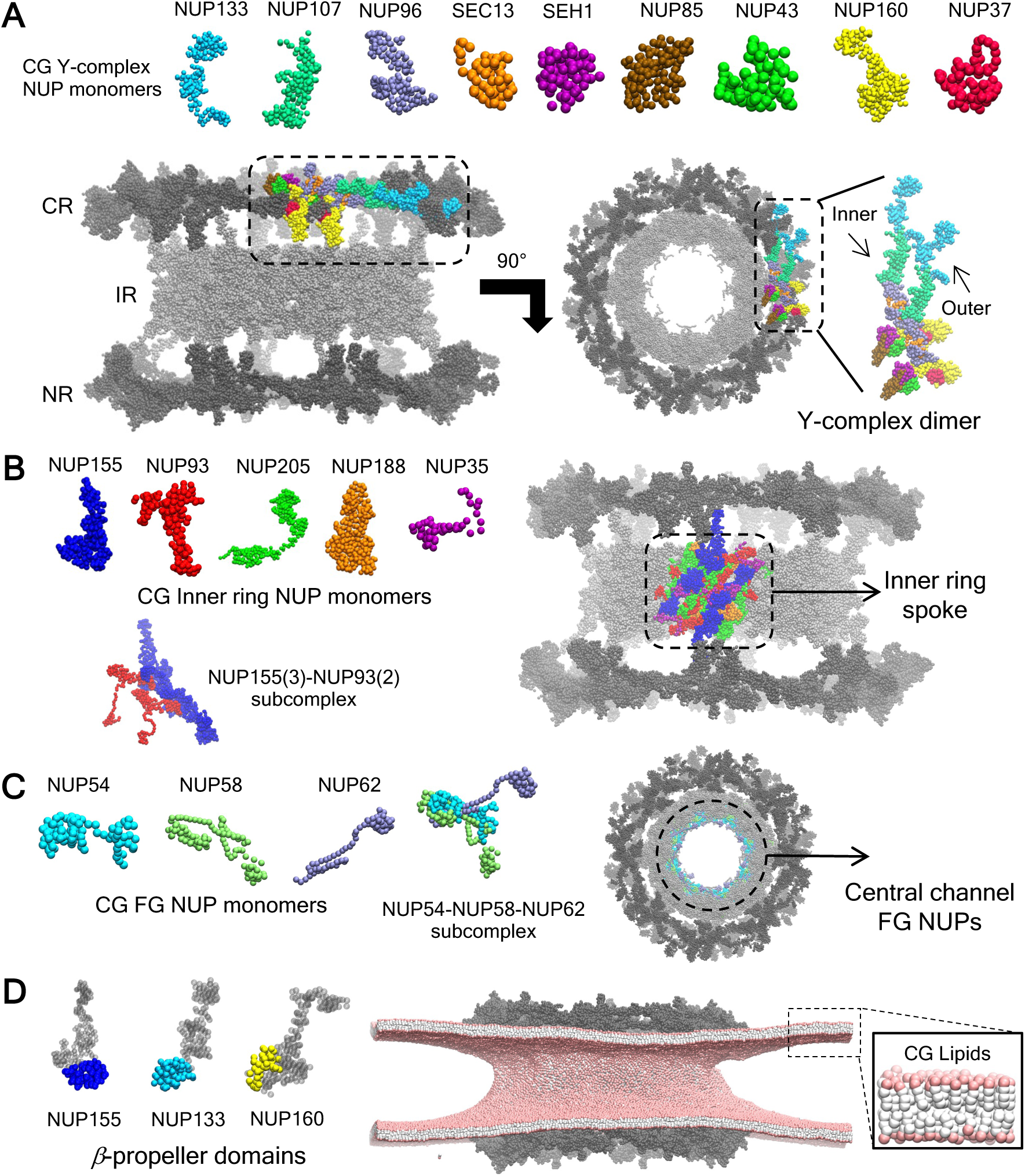
Overview of the CG molecular model of the CR, NR, and IR. (*A*) CG molecular model of each NUP monomer in the Y-complex in the upper panel. A side and top view of the NPC highlighting a single copy of the Y-complex dimer is shown in the lower panel. The rest of the NPC is shown in gray spheres. The inner and outer Y-complex monomer is labeled in the Y-complex dimer. (*B*) CG molecular model of each NUP monomer in the inner ring is shown in the upper left panel. The NUP155-NUP93 subcomplex consisting of 3 copies of NUP155 and 2 copies of NUP93 is shown in the lower left panel. A single unit of the inner ring spoke is shown (right panel) in the composite CG model of the human NPC and is highlighted in color. (*C*) CG molecular model of each FG NUP monomer and the NUP54-NUP58-NUP62 heterotrimeric subcomplex is depicted. A top view of the NPC is shown, highlighting the FG nucleoporins NUP62 lining the central channel. (*D*) In the left panel, the membrane binding *β*-propeller domains of NUP160, NUP133, and NUP155 are highlighted. The rest of the protein CG beads (no attractive interactions with lipids) are shown as gray spheres. In the right panel, the membrane-embedded composite dilated NPC model is shown. In the inset, the 4-site CG lipids are depicted. The headgroup of the CG lipid is shown in pink spheres. The interfacial CG bead and two tail beads of the CG lipid are shown in white spheres.

The CR and NR each consisted of 8 copies of the dimerized Y-complex (**Fig. 1*A***). The CG Y-complex consists of NUP133, NUP107, NUP96, SEC13, SEH1, NUP85, NUP43, NUP160, NUP37. Relative positions of the monomeric NUPs within a CG Y-complex and dimerization between the neighboring outer and inner CG Y-complex were maintained through a bonded soft elastic network model to allow configurational flexibility. Specifically, dimerization between an outer and inner Y-complex neighbor was maintained at the NUP133-CTD (inner) - NUP133-CTD (outer), SEC13 (inner) - NUP107-NTD (outer), and NUP160 (inner) - NUP96 (outer) interfaces. The head-to-tail oligomerization of the CG Y-complex dimers was modeled using non-bonded attractive interactions between a subset of CG sites at the NUP160-NUP133, NUP133-NUP37, and NUP107-NUP43 binding interface (**Fig. S1** of *Supporting Information*). The IR and CR/NR associations were modeled with attractive interactions between a subset of CG sites at the NUP155-NUP160 binding interface (**Fig. S2**). In both cases, the subset of CG site pairs with attractive interactions at the binding interface was determined from the corresponding atomic residues involved in direct contacts in the AA trajectories.

The CG model of the inner ring complex consisted of NUP155, NUP93, NUP205, and NUP188 (**Fig. 1*B***). The main building block of the CG inner ring spoke is a subcomplex composed of 3 adjacent copies of NUP155 and 2 copies of NUP93. Relative positions of monomers within this subcomplex were maintained with a bonded soft elastic network model. Interprotein interactions between NUP205 and NUP188 monomers and the NU155-NUP93 subcomplex within a spoke and between spokes were modeled with non-bonded attractive interactions (**Fig. S3**). Additionally, a polymer model of the NUP35 linker was used to connect the spokes. Phenylalanine-glycine (FG) containing channel nucleoporin NUP62 was modeled as a heterotrimeric subcomplex along with NUP54 and NUP58 (**Fig. 1*C***). The NUP54-NUP58-NUP62 subcomplex (**Fig. S4**) included a single copy of each protein, and each component within the subcomplex was connected through a bonded soft elastic network model. A total of 32 copies of NUP54-NUP58-NUP62 were anchored to the inner ring scaffold through non-bonded attractive interactions between NUP62 and NUP93 (NTD).

The nuclear membrane in our simulations was modeled by a 4-site CG lipid model with a bending rigidity of 30 *k_B_T* (38). We modeled the membrane association of the NPC through attractive interactions between the CG lipid head group and CG sites of the *β*-propeller domains of NUP160, NUP133, and NUP155 (**Fig. 1*D*** and **Fig. S5**). In our CG model, the minimum protein-membrane interaction strength was chosen for which all the *β*-propeller domains remained associated with the membrane. We then simulated the composite membrane-embedded constricted and dilated NPC model for 300 × 10^6^ CG MD timesteps (τ_*CG*_ = 50 fs). From the final 150 × 10^6^ *τ*_*CG*_ timesteps we estimated the diameter of the CR, NR, and IR. The mean diameters of the composite constricted and dilated CG NPC models closely reproduced the reference cryo-ET values (20), demonstrating the fidelity of our CG models (**Fig. S6**).

### Coarse-grained simulations of HIV-1 capsid translocation into the NPC central channel

To simulate HIV-1 capsid translocation into the NPC central channel, we derived a “bottom-up” solvent-free CG molecular model of the HIV-1 CA protein (46 CG sites), CA-CA attractive interactions to model full mature HIV-1 capsids, and attractive interactions between FG repeat (represented as a single CG site) and CA to model capsid binding to FG-NUPs. The reference all-atom (AA) MD trajectory from which the CG interactions were derived is a 3 CA hexamer complex with three 11-residue FG peptides, each bound to a CA hexamer (**Fig. 2*A***). A 1 microsecond long AA MD simulation trajectory was used to derive a CG molecular model of CA monomer (mapping resolution of ∼ 5 amino acids per CG bead). The AA trajectory was then mapped to a CG resolution. The CG non-bonded attractive interactions between CA monomers were derived from the mapped CG trajectory. We mapped the FG peptide to a single CG site to derive the CG non-bonded attractive interactions between the FG site and CA. We note that the FG-motif binding pocket is conserved across several host factors in the nuclear pore complex (39). In this work, we consider that the FG motifs in the N-terminal region (residue 1 to 150) of NUP62 also utilize the same binding pocket. Additional details of the HIV-1 capsid CG model are provided in the *Methods*.

**Fig. 2.**
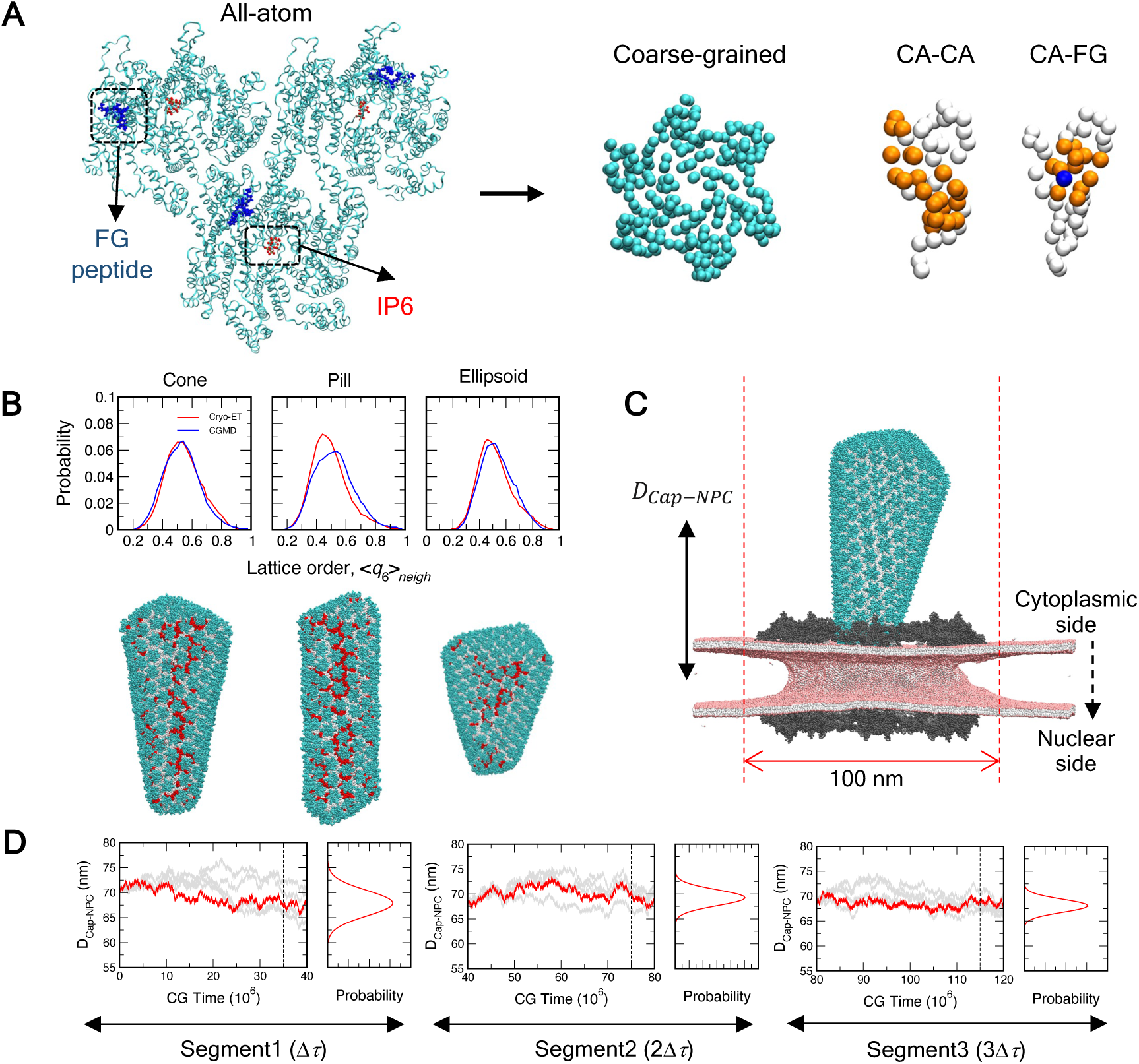
Schematic of coarse-grained (CG) HIV-1 CA model and CG MD simulation results. (*A*) A representative snapshot of the AA 18-mer CA complex (cyan cartoon) complexed with 3 FG peptides (residues SPSGVFTFGAN, shown in blue spheres) is shown in the left panel. The top right panel shows the representative snapshot of a CG CA hexamer (cyan spheres). The top right-most right panel shows (as orange spheres) the CG sites of a CA monomer involved in CA-CA and CA-FG associative interactions. The CG sites not involved in associative interactions in each case are shown as white spheres. The CG FG site associated with the CA monomer is shown as a blue sphere. (*B*) Comparison of the Steinhardt’s neighbor-averaged local bond order parameter (〈*q*_6_〉_*neigh*_) of HIV-1 capsid for the initial cryo-ET structure and from CG MD simulations (upper panel). The probability distribution of 〈*q*_6_〉_*neigh*_ for the CG simulations is calculated from the final 100 × 10^6^ CG MD timesteps, *τ*_*CG*_. The cone, pill, and ellipsoid capsids seen in the lower left panel of 2B contain 1314, 1122, and 1266 CA monomers, respectively. The lower left panel of 2B also depicts lattice disorder of the capsid lattice of the cone, pill, and ellipsoid morphology at the end of 400 × 10^6^ *τ*_*CG*_ of CG simulations (labeled as CG MD). The CTD domain of the CA monomers with 〈*q*_6_〉_*neigh*_ < 0.4 is colored in red. The CTD domain of the rest of the CA monomers is colored in white. The NTD domain of all the CA monomers is represented as cyan spheres. (*C*) A representative snapshot of an initial CG MD simulation configuration (side view) of the conical capsid placed at the cytoplasmic end of the NPC with the narrow end pointing to the central channel. The distance (*D*_*Cap-NPC*_) between the geometric center of the capsid and NPC inner ring is labeled. The dotted red lines indicate the location of cylindrical confinement of 100 nm diameter. The cytoplasmic and nuclear sides of the NPC are also labeled. (*D*) Schematic of the path sampling methodology to simulate translocation dynamics of HIV-1 capsid. For each segment, 5 parallel simulations (each 40 × 10^6^ *τ*_*CG*_) were evolved. From the probability distribution of *D*_*Cap-NPC*_ (calculated from final 5 × 10^6^ *τ*_*CG*_), the simulation trajectory was selected for which the endpoint (solid red line) is closest to the mean of the distribution. The endpoint of this trajectory was then used as the initial configuration for simulations in the next segment. The selected trajectories were appended to create the final translocation trajectory. The depicted segments correspond to the first 3 segments of the translocation trajectory of the HIV-1 capsid cone (narrow end pointing to the central channel) at the dilated NPC.

The initial atomic structure of the HIV-1 capsids was derived from cryo-ET images of capsids in intact virions and then mapped into CG structures (8, 40). To validate the CA-CA CG non-bonded associative interactions and whether these parameters maintain the capsid structural order, we simulated three different capsid morphologies (cone, pill, and ellipsoid) for 400 × 10^6^ CG MD timesteps, *τ*_*CG*_ (40). We characterized the capsid lattice order using neighbor-averaged Steinhardt’s local bond order parameter (〈*q*_6_〉_*neigh*_) (41, 42) for each CA monomer (see *Method* for details). Briefly, for a specific CA monomer, the higher the value of 〈*q*_6_〉_*neigh*_, the lattice contiguous to the CA monomer is more ordered. Conversely, the lower values of 〈*q*_6_〉_*neigh*_ indicate perturbation from the ideal lattice packing and therefore indicate more disorder and a likely weakening of the capsid structural integrity. The comparison of the distribution of the 〈*q*_6_〉_*neigh*_ values for each CA monomer calculated from the CG simulations, and initial cryo-ET configuration for all capsid shapes simulated are shown in **Fig. 2*B***. The final snapshot for each capsid depicts the lattice disorder classified as CA monomers with 〈*q*_6_〉_*neigh*_ < 0.4. The depicted domains in **Fig. 2*B*** are the capsid lattice’s intrinsic disorder (8). The 〈*q*_6_〉_*neigh*_ values calculated from the CG MD simulations closely match that of the initial configuration mapped from the cryo-ET images, therefore, helping to further validate the accuracy of the CA-CA CG interactions used to model the HIV-1 capsid in this work.

Using the aforementioned CG models of HIV-1 capsid and membrane-embedded human NPC, we then performed CG MD simulations of HIV-1 capsid translocation into the NPC central channel. All capsid translocation simulations are performed with cylindrical confinement of 100 nm diameter (**Fig. 2*C***) to restrict the diffusion of the capsid in the region coaxial to the central channel. The cylindrical confinement also emulates the cytoplasmic filaments (not included in the current CG NPC models) that likely play a role in directing the capsid to the NPC central channel. To perform the capsid translocation simulations into the NPC central channel, we implement a simulation protocol that is similar to parallel cascade MD (43). The simulation protocol used here allows for generating an effective trajectory that represents the likely pathway of the HIV-1 capsid translocation instead of performing multiple replicate simulations of capsid translocation thereby reducing the overall computational expense. The translocation of the capsids is tracked using a distance metric (*D*_*Cap-NPC*_), defined as the difference between the geometric center of the capsid and NPC inner ring along the direction of translocation (**Fig. 2*C***). A schematic overview of the simulation protocol is shown in **Fig. 2*D***. Complete details on the CGMD simulations of HIV-1 capsid translocation are provided in *Methods*.

### Capsid shape, the orientation of the approach, and channel dimension determine successful translocation events

To examine the factors that regulate the translocation of the viral capsids across the NPC central channel, we simulated the translocation dynamics of three different capsid morphologies (cone, pill, and ellipsoid), all with both the dilated and the constricted state of the NPC. For the cone-shaped capsid, we simulated two scenarios for which, in the initial configuration, either the narrow end or wide end was pointing to the NPC central channel. Overall, we generated 8 distinct capsid translocation dynamics trajectories from our simulations. The cumulative simulation time in these capsid translocation trajectories ranged between 400 to 1200 × 10^6^ *τ*_*CG*_. An attempted translocation is deemed unsuccessful if, within 400 × 10^6^ *τ*_*CG*_, the tip of the viral capsid proximal to the NPC fails to associate with the FG-NUPs of the central channel, and therefore, the simulations were terminated. For the trajectories in which the capsid successfully binds to the FG-NUPs, the simulations were extended for an additional 200 to 800 × 10^6^ *τ*_*CG*_. In **Fig. 3**, we show the time-series statistics of the distance (*D*_*Cap-NPC*_) between the geometric center of the capsid and equatorial midplane of the NPC inner ring along the channel axis for the successful capsid translocation simulations in. Additionally, we calculated the number of FG sites directly in contact with the CA monomers from the simulation trajectories to characterize the association of the capsid to the FG-NUPs at the central channel (shown in **Fig. 3** and **Supporting Movie 1 and 2**). Time-series statistics of the remainder of the translocation dynamics trajectories are shown in **Fig. S7-S9**. In **Fig. 4**, we show the final configuration of all 8 distinct capsid translocation dynamics simulations performed in this study.

**Fig. 3.**
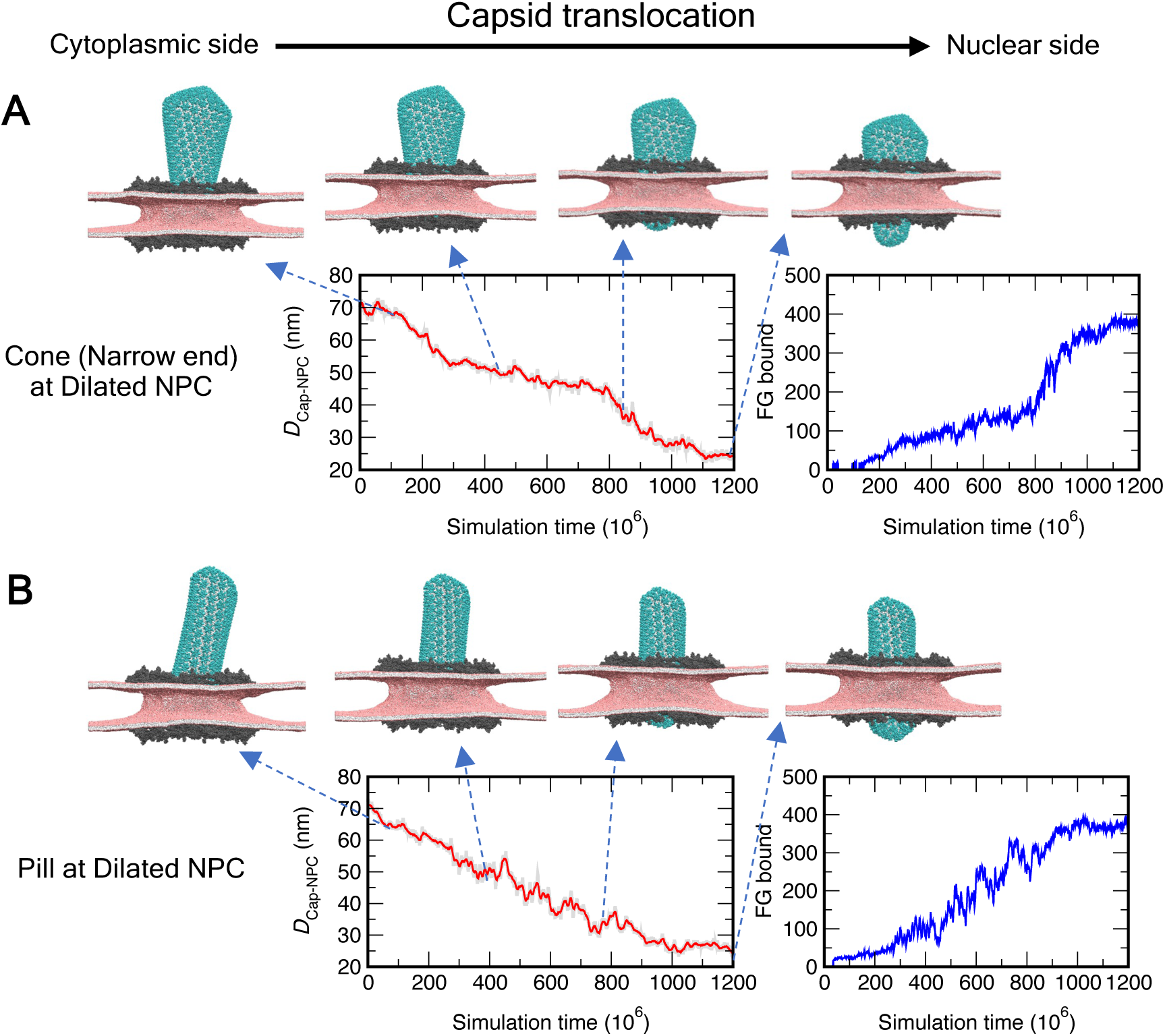
Translocation dynamics of HIV-1 capsid into the NPC central channel. Translocation dynamics time series plot (A and B) that depicts the distance (*D*_*Cap-NPC*_) between the geometric center of the capsid and NPC inner ring (left panel in red), and the number of FG sites of NUP62 bound to CA (right panel in blue). The snapshots depict the translocation of the cone-shaped (approaching from the narrow end) and pill-shaped capsid into the central channel of the dilated NPC from the cytoplasmic to the nuclear side at different points of the translocation dynamics trajectory. The capsid, NPC, and lipids are shown in the color scheme as in Fig. 2C.

**Fig. 4.**
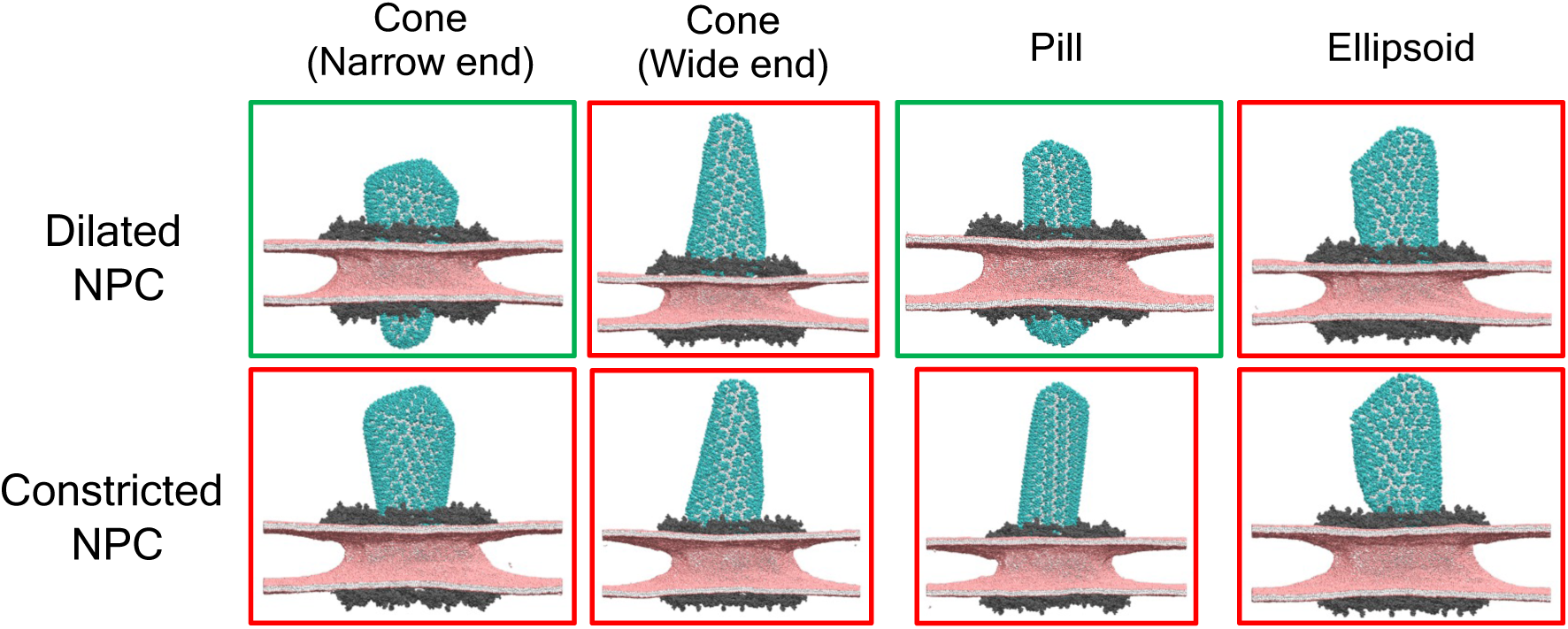
Configurations from the endpoint of the translocation dynamics trajectory for all capsid shapes and orientations. A green bordered panel shows the endpoint of a *successful* translocation trajectory in which the tip of the capsid translocates to the nuclear end. In our simulations, the cone-shaped capsid (when approaching from the narrow end) and pill-shaped capsid at the central channel of the dilated NPC successfully translocate to the nuclear end. Endpoints of the rest of the trajectories are shown in red bordered panels. The capsid, NPC, and lipids are shown in the color scheme as in Fig. 2C.

Our CG MD simulations reveal the behavior of different stages of viral capsid translocation facilitated by the FG-NUPs of the central channel. The cone-shaped capsid (approaching from the narrow end) and pill-shaped capsid at the dilated NPC form significant contacts with the central channel FG-NUPs within 400 × 10^6^ *τ*_*CG*_. In contrast, both the cone-shaped capsid (approaching from the narrow end) and pill-shaped capsid only have nominal contacts with the central channel FG-NUPs within 400 × 10^6^ *τ*_*CG*_ at the constricted NPC. The cone-shaped capsid (approaching from the wide end) and the ellipsoid capsid either only nominally associate or completely fail to associate with the central channel FG-NUPs for both the constricted and dilated state of the NPC. The early stages of the unsuccessful translocation dynamics trajectories reveal prolonged diffusion of the viral capsids at the cytoplasmic end of the NPC. However, the capsid tip in these unsuccessful translocation dynamics trajectories fails to translocate into the central channel.

We can rationalize the initial stages of capsid docking and translocation behavior by comparing the diameter of the central channel and the capsids examined in this study. The diameter of the inner ring scaffold, defined as the distance (*D*_*NUP*188_) between the geometric center of the NUP188 at the inner ring of the opposite spokes, is 59 ± 7 nm and 73 ± 6 nm for the constricted and dilated states of the NPC, respectively. The capsid binding FG-NUPs lining the central channel are associated with the inner ring scaffold. Here, we approximate the void cross-section of the central channel as the distance between the geometric center of the NUP62 of the opposite spokes (**Fig. 1C**). The void cross-section of the central channel available for the capsid to occupy is 41 ± 4 nm and 56 ± 6 nm for the constricted and dilated state of the NPC, respectively. The diameter of the conical capsid at the narrow and wide end is 33 and 59 nm, respectively. The diameter of the pill and ellipsoid capsids are 42 and 56 nm, respectively. The diameter of the narrow end of the capsid cone is considerably smaller than the void cross-section of both the constricted and dilated states of the NPC; hence the cone can spontaneously associate with the central channel of both constricted and dilated NPC in our simulations. The size compatibility also explains the association of the pill-shaped capsid to the dilated state of the NPC. After initial stable association to the central channel FG-NUPs is established, the cone-shaped and pill-shaped capsid at the dilated NPC continuously translocates from the cytoplasmic to the nuclear end.

The time-series profiles of translocation from the cytoplasmic to the nuclear side of the NPC reveal key molecular insight into the factors that regulate successful passage. The cone initially undergoes a period of rapid passage through the central channel (*τ* < 300 × 10^6^ *τ*_*CG*_), followed by a period during which the rate of translocation progressively slows down (300 × 10^6^ < *τ* < 800 × 10^6^ *τ*_*CG*_). In the final stages (800 × 10^6^ < *τ* < 1100 × 10^6^ *τ*_*CG*_), the capsid again undergoes rapid passage. It is interesting to note that until *τ* < 800 × 10^6^ *τ*_*CG*_ the number of FG sites in contact with CA slowly and monotonically increases. After that, there is a sharp increase in the number of FG sites in contact with CA. We, therefore, propose that the translocation of the cone-shaped capsid adheres to the following successive step: During the initial period of rapid translocation, the narrow end binds to the FG-NUPs of the central channel. The efficiency of passage gradually decreases as the central channel encounters the wider regions of the capsid. This can be attributed to the increasing steric interactions between the capsid and non-FG-NUPs of the central channel. In between 800 × 10^6^ < *τ* < 1100 × 10^6^ *τ*_*CG*_, the steep increase in the binding of FG sites to CA appears to facilitate rapid passage of the capsid cone. It also appears that at this stage, the adhesion between inner ring NUPs is loosened from the mechanical stress applied from the capsid allowing structural relaxation of the central channel. The consequent dilation of the channel assists in the accommodation of the wider regions of the cone-shaped capsid. In contrast, the pill-shaped capsid translocates into the central channel at a constant rate which can be attributed to its uniform cross-section.

To summarize, our extensive bottom-up CG MD results suggest that successful viral capsid passage across the NPC central channel is facilitated by the energetics of the capsid-binding FG-NUPs, while the primary barrier to translocation originates from the incompatibility between the channel dimension and capsid morphology.

### Distinct lattice disorder patterns are formed in intact HIV-1 capsids docked at the NPC central channel

Visual inspection of the capsid structures at the endpoint of the translocation trajectories revealed that cone-shaped and pill-shaped capsids remained intact. To assess the stress impact of the spatially confined environment of the NPC central channel on the capsid structural state, we characterized the capsid lattice order using neighbor-averaged Steinhardt’s local bond order parameter (〈*q*_6_〉_*neigh*_) (41) per CA monomer for the cone-shaped and pill-shaped capsids docked at the dilated NPC. We considered the capsid configurations at different stages of translocation at the NPC central channel (**Fig. 3**). To quantify the extent of lattice disorder, we identified the CA monomers with 〈*q*_6_〉_*neigh*_ < 0.4 and classified these monomers as disordered. We then characterized the size and organization of these disordered domains at different stages of translocation of the capsid into the central channel by performing clustering analysis and finally calculated the radius of gyration (*R*_*g*_) of the largest disordered cluster. To perform the analyses, we divided the 1200 × 10^6^ *τ*_*CG*_ long CG MD translocation trajectory of cone and pill-shaped capsid at the dilated NPC into 4 equal segments and label the segments as Stage 1-4.

Isolated uncorrelated disordered domains are intrinsic to the capsid, observed both in the initial cryo-ET structure and in a free capsid in CG MD simulations (**Fig. 2*B***). As the capsid approaches and associates with the NPC central channel, we can identify distinct phases of weakening of the structural integrity of the viral capsid. **Figures 5*B*** and **5*C*** display the average size of the largest contiguous disordered domain of CA monomers and the radius of gyration (*R*_*g*_) at different stages of capsid passage. Initially, the disordered domains of the capsid grow independently of each other. The initial stages (1 and 2) encompass the approach of the capsid to the NPC central channel at the cytoplasmic end to the first stable association to the FG-NUPs of the central channel. In the later stages (3 and 4), the capsid translocates into the central channel driven by binding to the FG-NUPs while experiencing spatial confinement. In the later stages, we observe a significant increase in the disordered domain size consisting of greater than 20 CA monomers (**Fig. S10**). We attributed the growth of the disordered domains to the steric interactions of the capsid with the cytoplasmic ring of the NPC scaffold. The disordered domains are large enough in size to anneal into well-defined striated patterns on the capsid lattice (**Fig. 5** and **Fig. S11**). The largest disordered domain contains ∼ 60 CA monomers with a dimension up to ∼ 20 nm when the tip of the capsid reaches the nuclear end. These striated disordered patterns on the capsid lattice are quite dynamic and undergo continuous rearrangement in the timescales of our CG MD simulation. Whether thermal fluctuations can lead to the stochastic nucleation of cracks at these striated disordered patterns will likely require significantly longer simulation timescales.

**Fig. 5.**
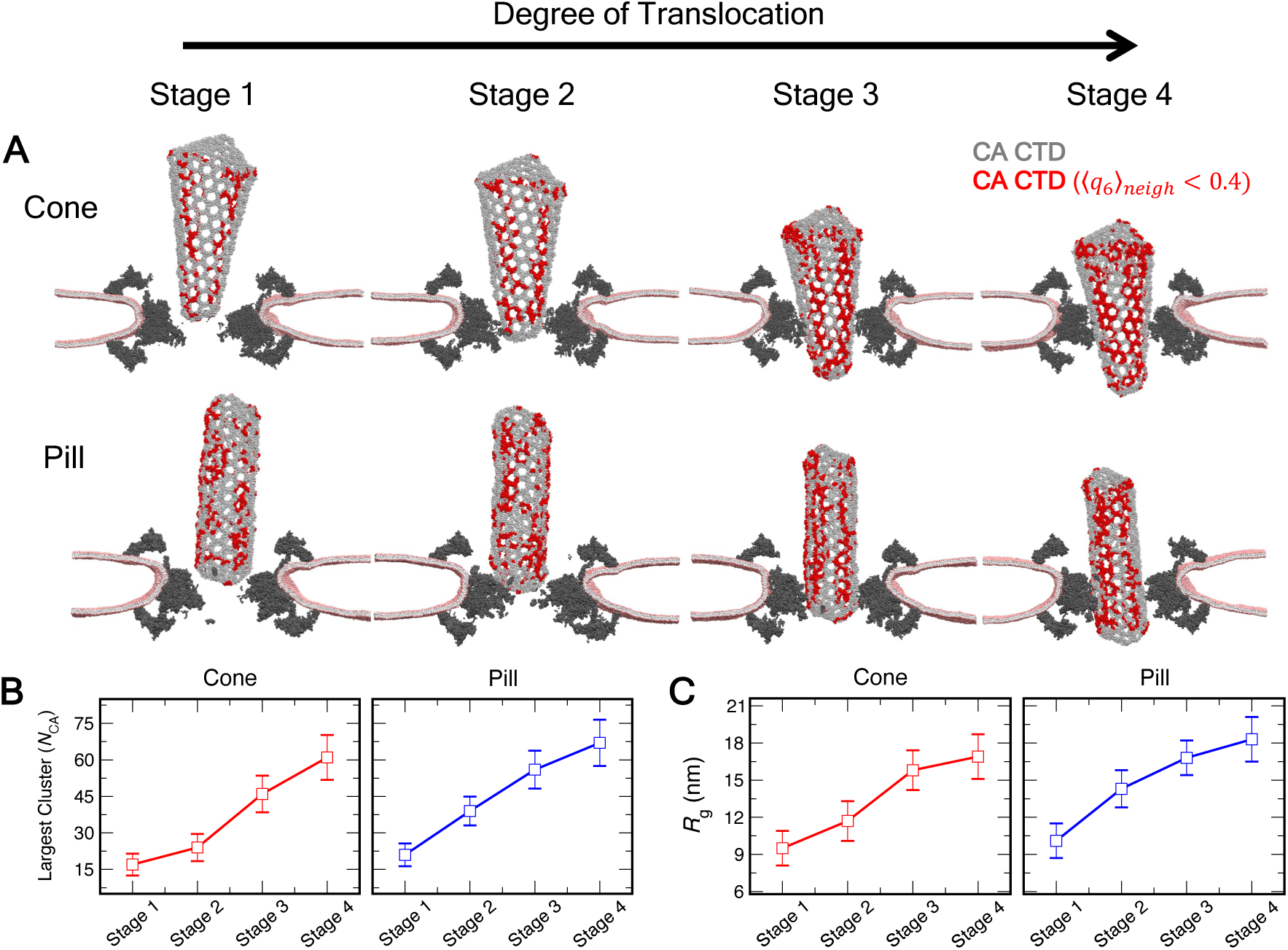
Lattice disorder analysis of HIV-1 capsid at different stages of entry into the NPC central channel. (*A*) The translocation dynamics trajectory of cone-shaped (approaching from the narrow end) is divided into 4 equal sections to define four stages of capsid docking to the NPC central channel. The identical protocol is used to define the 4 stages of entry for the pill-shaped capsid. In each snapshot (at the endpoint of each stage), a cutaway sideview of the NPC (shown in gray spheres) and lipid bilayer is shown. The headgroup bead of the CG lipids is shown in pink spheres. The interfacial and tail beads are shown in white spheres. The CTD domain of capsid is shown in silver spheres. The CTD domain of each CA monomer with 〈*q*_6_〉_*neigh*_ < 0.4 is shown in red spheres. The CA NTD domain is not shown for clarity. The snapshots correspond to the endpoint of the simulation at each stage (300, 600, 900, and 1200 × 10^6^ *τ*_*CG*_). (*B*) Largest connected disordered cluster statistics of each stage for translocation of the cone-shaped and pill-shaped capsid. Here, *N*_*CA*_ is the number of CA monomers that constitute the largest connected disordered cluster in our analysis. (*C*) The radius of gyration (*R*_*g*_) of the largest connected disordered cluster at each stage for translocation of cone-shaped and pill-shaped capsid into the central channel.

At this point, it is tempting to conclude that the CA protein lattice disorder of the capsid seen upon its translocation into the NPC is a measure of an “injured” capsid, that may in turn affect its later uncoating behavior within the nucleus. Such a conclusion indeed seems justified. We note that when docked at the NPC central channel, in addition to the energetically favorable FG-NUP interactions, the capsid also encounters non-FG-NUPs at a significantly higher frequency. The latter is energetically unfavorable and contributes to stress of the capsid lattice. The ability of the capsid lattice to become more disordered upon experiencing the stress from its interaction of the NPC may also play a dual role. This disordering behavior is a sort of “plasticity” so that the capsid can better absorb the stress from the NPC interaction. Such behavior is often referred to as an “entropic spring” in which the increase in entropy (lattice disorder, i.e., Δ*S* > 0) means the capsid lattice free energy is lowered to a degree in response to the stress (−*T*Δ*S* < 0). Without that plasticity, the capsid CA lattice would be very fragile and more easily crack prematurely (in terms of the virial replication lifecycle), while within the NPC.

### Condensation state of RNP modulates the lattice disorder of HIV-1 capsids

Recent studies have shown how the RNP, and eventual reverse transcription of the viral genomic complex appears to impact the structural integrity of the capsid (40, 44). Mature capsids encapsulate ribonucleoprotein (RNP) complexes in a condensed globular form during viral assembly (45). Reverse transcription leads to uncoating and subsequent stiffening of the viral genomic RNA. The increased spatial requirement of the genomic complex results in the loss of capsid patches allowing the extrusion of the newly synthesized DNA (18, 19, 44, 46). To simulate capsids docked at the NPC (corresponding to Stage 4 in **Fig. 5**) with additional biological detail, we incorporated two polymeric chains in the capsid interior that minimally emulate two 9000-nt RNP complexes. Briefly, modulating the interaction strength (*ε*_*RNP*_) between the CG beads of the polymer allows for controlling the condensation state of the RNP complex in our simulations. Additionally, CG beads of the RNP weakly interact with the C-terminal domain (CTD) tail of CA. Full details of the RNP model are provided in the *Methods*. We simulate the RNP complex in the capsid interior by varying *ε*_*RNP*_ between 0.3 and 0.9 kcal/mol. For each *ε*_*RNP*_ value simulations are evolved for 80 × 10^6^ *τ*_*CG*_. We perform 3 replica simulations for each *ε*_*RNP*_ for both the cone-shaped and pill-shaped capsid. Here, our goal is to establish how different condensation states of the RNP complex impact the structural integrity and elastic response of the capsid docked at the NPC. The condensation states of the RNP complex in our simulations are intended to model the viral genome at different stages of uncoating prior to or at the initiation of reverse transcription.

**Figures 6*A*** and **6*B*** summarize the results of the simulations of the RNP in the interior of the cone-shaped and pill-shaped capsid for different *ε*_*RNP*_ values. Interestingly, the degree of RNP condensation, as characterized by the radius of gyration (*R*_*g*_) of the polymer complex, appears to be sensitive to both *ε*_*RNP*_ and the shape of the capsid (**Fig. S12**). At *ε*_*RNP*_ = 0.9 kcal/mol, the RNP complex spontaneously self-associates and forms a globular condensate (**Fig. S13**). For the cone-shaped capsid, the globular RNP complex localizes at the wide end in our simulations. Our simulations recapitulate the observation of imaging studies of preferential localization of the RNP complex at the wide end of the cone-shaped capsid (45, 47), thus further establishing the validity of the minimal CG RNP model used in our studies. We find that for the pill-shaped capsid, the RNP also condenses and localizes at the tip of the capsid. For both the capsids, we attribute the preferred localization of the condensed RNP complex at the capsid tip to maximizing the contacts between the C-terminal end of CA and RNP beads at the surface of the condensate. Interestingly, in all the replica simulations, the condensed RNP in the pill-shaped capsid localizes at the tip distal to the NPC central channel. We speculate that the free and unconfined tip of the pill-shaped capsid can undergo enhanced volume fluctuations which allows accommodation of the condensed RNP complex. At *ε*_*RNP*_= 0.6 kcal/mol, the RNP complex forms a partially condensed core with extended ends (**Fig. S13**). At *ε*_*RNP*_= 0.3 kcal/mol, the RNP complex becomes fully uncondensed, associating to C-terminal end of CA throughout the interior of the capsid (**Fig. S13**). For the pill-shaped capsid, at *ε*_*RNP*_ = 0.9 kcal/mol, localization of the fully condensed RNP complex at the tip generates localized cracks (**Fig. S14**). These cracks then lead to a degree of extrusion of the RNP in the uncondensed form (*ε*_*RNP*_= 0.3 kcal/mol). The appearance of these localized lattice cracks can be attributed to the significantly lower interior volume of the pill compared to the cone. We note that, despite the formation of localized cracks, the overall morphological integrity of the capsid is still preserved in our simulations, perhaps again reflecting a possible “entropic spring” behavior.

**Fig. 6.**
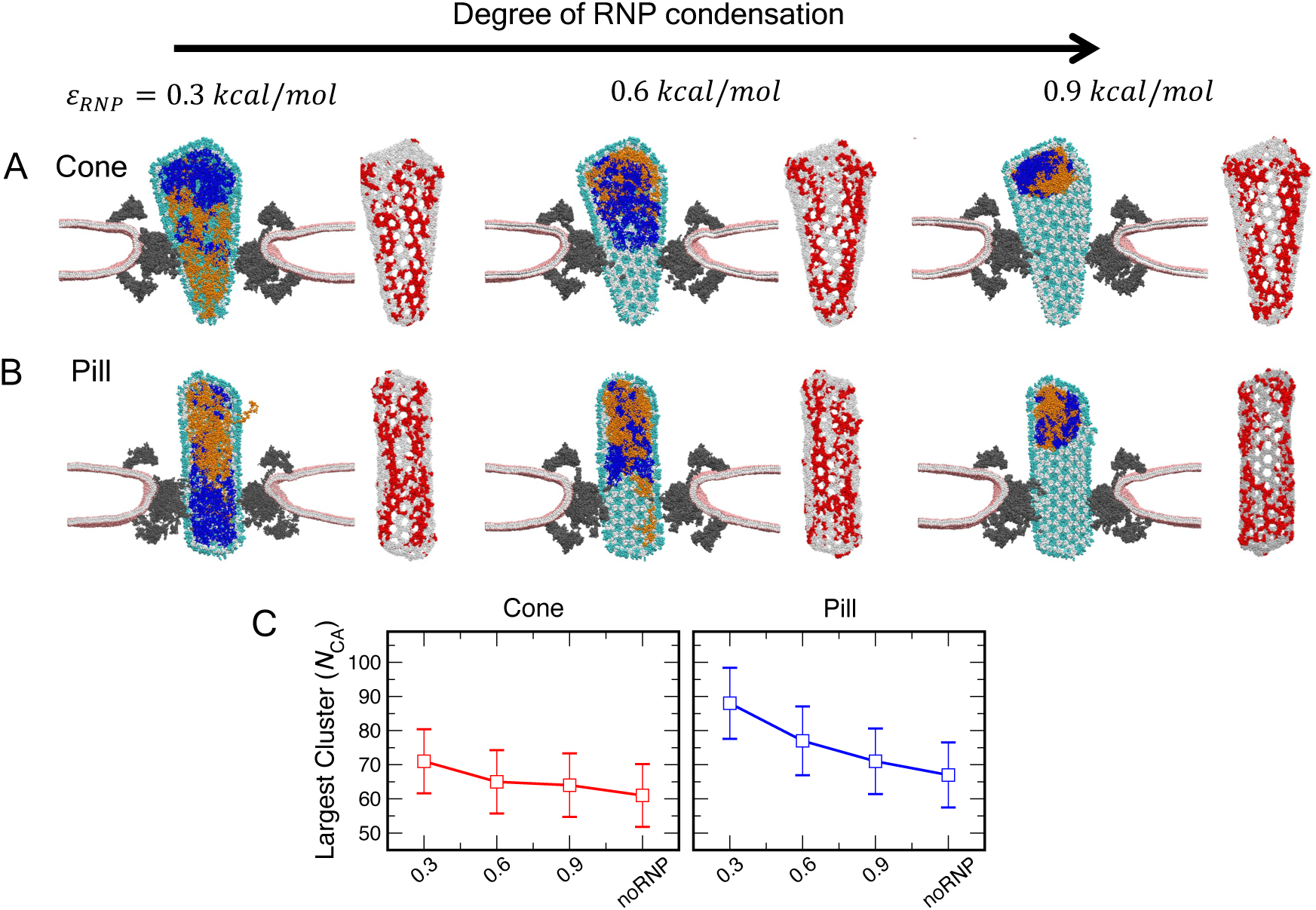
Spatial localization of the RNP complex in the capsid interior docked at the central channel and lattice disorder analysis. In each snapshot, a cutaway sideview of the NPC and lipid is shown with the same colors as in Fig. 5. The capsid lattice is vertically sliced to show the RNP complex in the interior. The two RNP chains are shown in blue and orange spheres. From left to right, *ε*_*RNP*_ is varied from 0.3 to 0.9 kcal/mol, thereby modulating the RNP condensation states. For both the cone (*A*) and pill (*B*), fully condensed RNP localizes at the capsid tip away from the NPC central channel. For each *ε*_*RNP*_, the capsid with lattice disorder patterns is also shown on the side. The CTD domain of capsid is shown in silver spheres. The CTD domain of each CA monomer with 〈*q*_6_〉_*neigh*_ < 0.4 is shown in red spheres. The CA NTD domain is not shown for clarity. The snapshots correspond to the endpoint of the simulation. (C) Largest connected disordered cluster statistics for each *ε*_*RNP*_ value simulated in this study for the cone and pill.

The condensation state of the RNP complex can create variable degree of internal mechanical stress on the capsid lattice in addition to external stress from the steric confinement arising from the central channel. To characterize the impact of the internal and external stress on the structural integrity of the capsid, we compute the lattice order of capsids docked at the NPC central channel at different RNP condensation states (**Fig. 6*C***). As seen in **Fig. 6*C***, the degree of RNP condensation has no significant effect on the largest disordered cluster for the cone. In contrast, there is a significant increase (∼ 50%) in the largest disordered cluster size for the pill compared to when there is no RNP in the capsid interior. Our results therefore appear to demonstrate that the structural integrity of the pill is significantly weakened compared to the cone due to the lower interior volume of the former. We speculate that while both the cone and pill can successfully dock at the NPC central channel, the lattice packing of the pill is significantly weakened, which can lead to partial disassembly of the capsid during nuclear entry.

### Progressive dilation of the NPC central channel facilitates capsid translocation

To this point, we have established that the dilated state of the NPC allows passage of the cone-shaped and pill-shaped capsid. We next sought to characterize how mechanical stress arising from the translocation of the viral capsid impacts the inner ring scaffold structural organization. It is known that the inner ring spokes are capable of sliding movements under mechanical stress while preserving the arrangement of constituent NUPs within the spokes (48, 49). To analyze the structural change to NPC upon capsid translocation, we measure the diameter of the central channel. The central channel diameter is defined as the distance between the geometric centers of NUP188 at the opposite spokes of IR (**Fig. 7**). Translocation of both cone-shaped (Fig. 7A) and the pill-shaped capsid (Fig. 7B) cause additional dilation of the IR scaffold. However, the degree of dilation as a consequence of translocation for the cone-shaped capsid (∼ 11%) is greater than the pill-shaped capsid (∼ 7%) when the tips of the capsid reach the nuclear end (Stage 4). The higher degree of dilation of the channel for the translocation of the cone compared to the pill reflects the difference in size between the capsids (59 nm at the wide end of the cone and 42 nm for the pill). Based on our analysis, it is evident that modest dilation of the IR scaffold is key to alleviating the steric stress and facilitating translocation of the capsid.

**Fig. 7.**
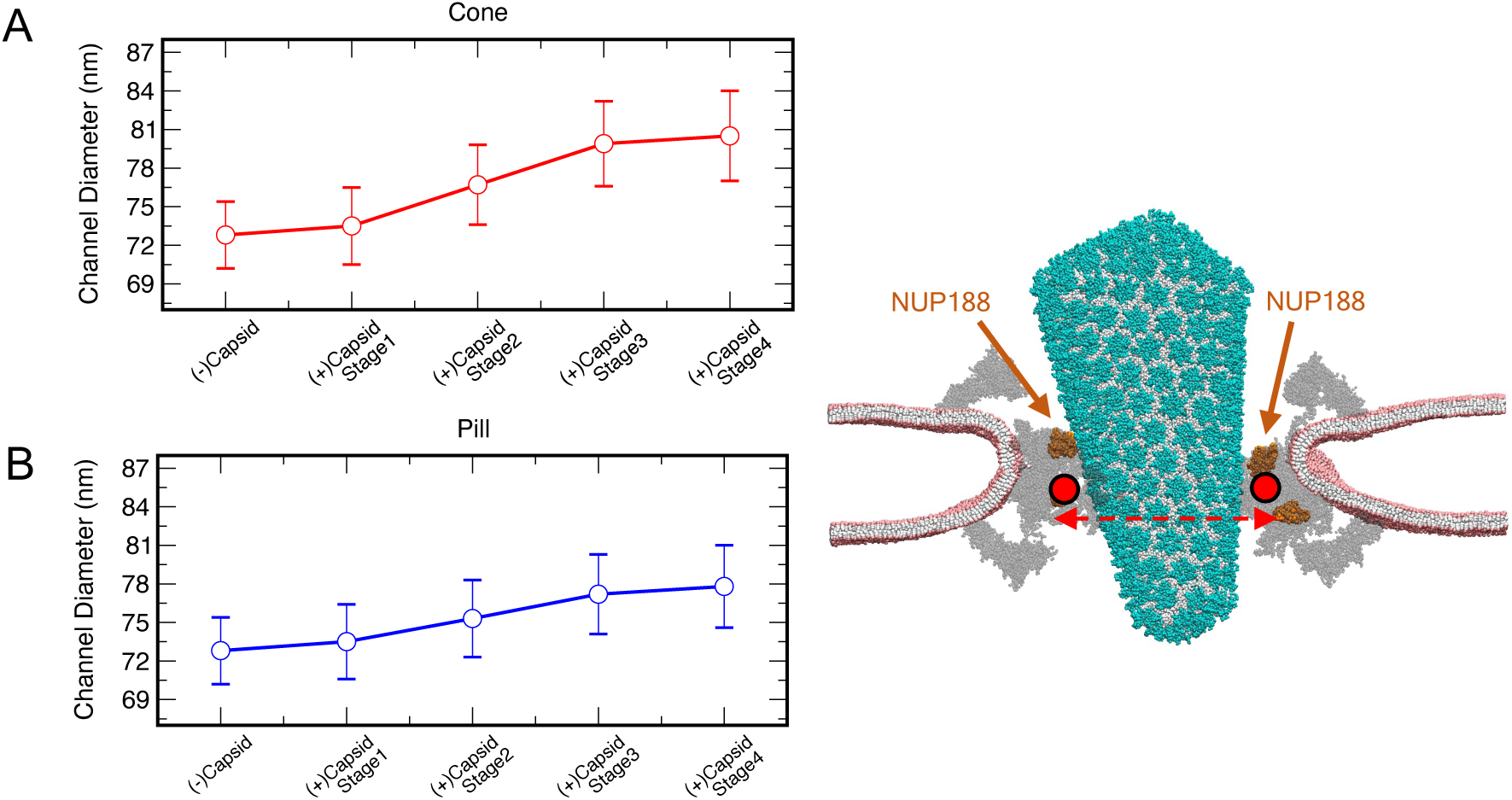
NPC central channel diameter at different stages of capsid docking. Measurement of the central channel diameter for the translocation simulation trajectories of the cone (A) and pill (B) into the dilated NPC. The channel diameter of the dilated NPC without capsid is calculated from the final 150 × 10^6^ *τ*_*CG*_ of a 300 × 10^6^ *τ*_*CG*_ CG MD trajectory. The data point (-)Capsid represents the channel diameter in the absence of the docked capsid. The data points (+)Capsid Stage 1-4 represent capsid entry at different stages (same as in Fig. 5 and 6). The data points report the mean and standard deviation calculated over the 300 × 10^6^ *τ*_*CG*_ for each stage of the capsid translocation simulation. The right panel shows the cutaway sideview of the NPC and lipid. The NUP188 of the IR is shown in orange spheres. The estimation of the channel diameter is performed between the geometric center of NUP188 of the opposite IR spokes (shown in horizontal red dashed line). The mean channel diameter for the reference cryo-ET structure of the dilated NPC is 70.4 nm.

### Cohesiveness of disordered NUP98 impedes capsid association to the central channel

The disordered FG-NUP98 is a component of the NPC central channel that can regulate nucleocytoplasmic passage of macromolecules (50, 51). Adjacent chains of NUP98 can engage in multivalent cohesive interactions creating a hydrogel-like environment that likely impacts the translocation dynamics of macromolecules (50, 52, 53). To investigate how the polymer-mesh characteristics of NUP98 chains grafted to the NPC can help to regulate capsid translocation, we built a CG polymer model of NUP98_1-620_ and attached 48 NUP98 chains to the dilated CG NPC model. We hereafter refer to this updated model as NPC_NUP98_. In the NPC_NUP98_ model, each NUP98 chain was anchored to NUP155 in the dilated NPC model based on cryo-ET structural data (20). In our simulations, we modulated the NUP98 interaction strength (*ε*_*NUP*98_) of the FG-rich regions between 0.2 and 1.0 kcal/mol. The CG beads of NUP98 containing FG-motif also have attractive interactions with CA, as described in the previous section. To simulate the association of the narrow end of the cone-shaped capsid, we performed CG simulations for 60 × 10^6^ *τ*_*CG*_ for each *ε*_*NUP*98_. We performed 3 replica simulations for each *ε*_*NUP*98_.

Figure 8 summarizes the results of different cohesive states of NUP98 and the association of the narrow end of the cone-shaped capsid to the NPC central channel. For the weaker *ε*_*NUP*98_ interactions, the NUP98 chains formed extended configurations creating a highly dynamic and interactive mesh-like environment and occupying the void space of the central channel. We note that *in vivo*, when attached to the NPC, the NUP98 chains adopt extended configurations (50). Therefore, the weaker *ε*_*NUP*98_ in our simulations most accurately mimics the disordered NUP98 conformations in the NPC environment. As the *ε*_*NUP*98_ is increased, the NUP98 chains become gradually cohesive, forming local condensates in the vicinity of the IR scaffold. For the weaker *ε*_*NUP*98_ interactions, as the tip of the capsid approaches the central channel, FG-motifs of the NUP98 chains contacts the capsid. The binding of the multiple NUP98 FG-motifs immerses the narrow end of the capsid in the unstructured mesh-like environment at the NPC central channel and promotes translocation of the capsid (**Fig. 8*A***). In contrast, for the stronger *ε*_*NUP*98_ interactions, the tip of the capsid encounters local condensates of NUP98 chains in which the FG-motifs bind the CA lattice with significantly reduced propensity compared to the weaker *ε*_*NUP*98_ interactions (**Fig. 8*B***). Therefore, the lack of FG-motif binding to the constituent CA monomers regulated by the NUP98 condensation state impedes the association of the narrow end of the cone-shaped capsid to the NPC central channel.

**Fig. 8.**
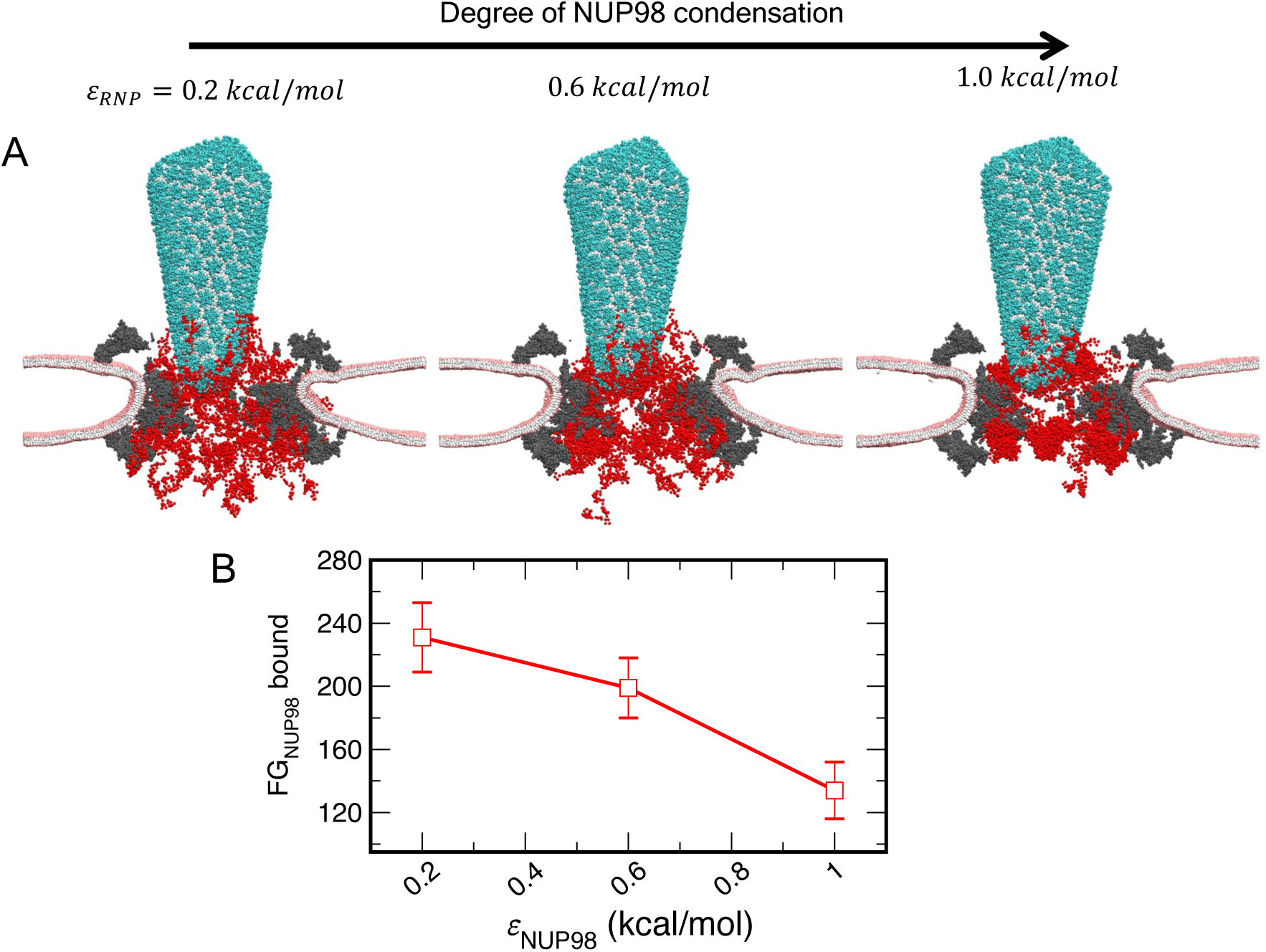
Condensation state of NUP98 at the NPC_NUP98_ central channel and capsid tip association. (A) In each snapshot, a cutaway sideview of the NPC and lipid is shown with the same colors as in Fig. 5. The NUP98 chains are shown in red spheres. From left to right, *ε*_*NUP*98_ is varied from 0.2 to 1.0 kcal/mol, modulating the NUP98 condensation state. For each snapshot, the association of the capsid tip to NUP98 mesh is shown. (B) The number of FG sites of NUP98 chains bound to CA. Note that each NUP98 consists of 12 FG-motif sites, and there are 48 NUP98 chains tethered to NPC_NUP98_. The standard deviation is calculated over 3 replica simulations for each *ε*_*NUP*98_.

The results of our simulations present the following mechanistic picture. To successfully associate to the NPC central channel, the capsid tip must permeate through the mesh-like medium of NUP98 chains and disrupt the inter-NUP98 interactions. Therefore, favorable FG-CA interactions must overcome the inter-NUP98 interactions that determine the cohesiveness of the mesh-like medium. As a result, capsid association with the NUP98 chains is only energetically feasible for weaker inter-NUP98 interaction strengths.

## DISCUSSION and CONCLUDING REMARKS

How the NPC allows selective and efficient transport of large cargoes remains an extremely important area of research. The mechanistic details of the nuclear import of the HIV-1 viral capsids, which have comparable dimensions to the NPC central channel, are yet to be fully understood. Importantly, the nuclear import step of the viral capsid can be a key target for small molecule capsid inhibitors (54, 55). In this study, we developed and utilized bottom-up CG MD simulations to elucidate the mechanisms that regulate the docking and translocation of the HIV-1 viral capsid into the central channel. Specifically, we explored how capsid morphology and orientation of approach regulate a successful translocation event. We observed successful translocation of conical and pill-shaped viral capsids for the dilated state of the NPC. Our simulations recapitulate recent visualization of cone-shaped and tube-shaped intact capsids through cryo-electron tomography (cryo-ET) and electron microscopy at the central channel of the NPC and nucleoplasm of infected and non-infected cell (18). Analysis of our CG MD translocation dynamics trajectories sheds light on the factors that are key to the nuclear import of viral capsids and how the effective elasticity and structural integrity of the capsid is impacted. We anticipate that the mechanistic insights obtained in this work can also be broadly applied to infer the pathway of import of other large cargoes across the central channel of NPC.

The cross-section of the entire conical capsid is compatible with the dimension of the dilated state of the HIV-1 capsid. One of our key findings is that despite the size compatibility of the entire capsid cone and NPC central channel in the dilated state, successful docking and translocation are contingent on the orientation of the capsid approach. Our simulations suggest that when approaching from the wide end and despite the size compatibility of the conical capsid with the NPC central channel, the initial steric barrier is too high to be surmounted by energetic contributions from interactions of CA with FG-NUPs at the central channel (56). Furthermore, at the wide end of the capsid cone, there are a higher number of available CA proteins for the channel FG-NUPs to bind relative to the narrow end, therefore providing a thermodynamic driving force for the association of the capsid cone at the wide end. Hence, the inability of the cone-shaped capsid in that orientation to translocate into the central channel is likely due to a kinetic barrier. When approaching from the narrow end, the cone-shaped capsid associates at the cytoplasmic end of the NPC central channel and translocate till the tip of the capsid reaches the nuclear end. Importantly, the translocation of the cone-shaped capsid continuously dilates the IR scaffold to progressively accommodate the wider regions of the cone. The preferred mechanism of the translocation of the capsid approaching from the narrow end is consistent with *in situ* cryo-ET image of the HIV-1 core and capsid docking to NPC mimicking DNA-origami-based channel (18, 57). The binding of the capsid to the central channel generates outward stress. Therefore, progressive dilation of the central channel is key to increasing the effective void volume to accommodate a non-uniform shape cargo like the conical capsid and reduce steric interactions. The IR scaffold architecture appears to be malleable enough to weaken the interactions between the constituent NUPs and allow dilation of the central channel under conditions of mechanical stress. Therefore, future experimental and enhanced sampling atomistic simulation studies are warranted to understand better how different NUP subcomplexes at the IR scaffold respond to mechanical stress and trigger large-scale conformational changes of the NPC (49).

We also analyzed how the capsid structural state is perturbed when translocating into the NPC central channel. Uncorrelated small, disordered domains are intrinsic to the capsid lattice. We find that when docked to the NPC central channel, spatial confinement and steric interactions with central channel NUPs augment the number and the size of the disordered domains at the capsid lattice. Importantly, these isolated disordered domains anneal to highly correlated striated patterns on the capsid lattice. The presence of an uncondensed RNP complex in the capsid interior further amplifies the size of the correlated disordered domains on the capsid lattice due to internal stress. The effect of the fully uncondensed RNP on capsid lattice disorder is particularly significant in the case of the pill-shaped capsid due to the lower interior volume when compared to the cone-shaped capsid. Importantly, both cone-shaped and pill-shaped capsid remains morphologically intact during translocation in the timescales of the CG MD simulations. Our results are consistent with the notion that primarily intact capsids enter through the NPC central channel (18). Finally, our analysis indicates that while intact capsids dock at the NPC central channel, the cone, in contrast to the pill, will most likely translocate to the nuclear side without significant weakening of the structural integrity.

The observation of the distinct disordered patterns on the capsid lattice in our simulations docked at the NPC indicates a significant weakening of the structural integrity of the capsid (40), but at the same time a certain overall capsid “plastic” response to the NPC stress. It is known that nucleation of the first defect is the trigger for capsid rupture, while disassembly of capsomers progressively occurs post lattice rupture (58). Additionally, the initiation of reverse transcription can also impact the capsid structural stability (46). The highly correlated striated patterns observed at the capsid lattice when docked at the NPC central channel in our simulations can be the birthplace of defects remodeling the capsid shape during the nuclear entry (18).

As the capsid tip reaches the nuclear side of the NPC, it will encounter capsid binding NUP153 in the NPC nuclear basket (59). A detailed structure of the nuclear basket is yet to be established and hence is not yet included in our CG NPC model. NUP153 plays a key role in regulating the late stages of capsid nuclear entry (25, 60). Multiple recent studies have elucidated how NUP153 associates with HIV-1 capsid (61-63). It is noteworthy that NUP153 preferentially targets the high curvature regions of the capsid lattice (63). Whether encountering NUP153 at high densities at the nuclear basket stabilizes the structurally fragile capsid after passage through the NPC central channel will require additional studies.

Given the complexity of the human NPC, our current CG NPC model consists of CR, NR, and IR. The human NPC also consists of the eight cytoplasmic filaments and capsid binding FG-NUPs (NUP358, NUP214, and NUP88) localized at the cytoplasmic side (21, 64). The cytoplasmic filaments and the FG-NUPs at the cytoplasmic side likely play an important role in orienting the capsid for translocation into the central channel. The cytoplasmic filaments and the cytoplasmic FG-NUPs are not included explicitly in our current model. Thus, the initial configuration of the capsids in our simulations corresponds to the sequence of events after the FG-NUPs at the cytoplasmic side position the tip of the capsid to create optimal circumstances for interaction with central channel FG-NUPs. Our CG NPC model is also designed to represent the discrete constricted and dilated states. NPC is pleomorphic and explores multiple intermediate states between the constricted and dilated states under varying mechanical conditions triggered by conformational change of specific NUP subcomplexes (49, 65) and variation of nuclear membrane mechanical properties (56, 57). One can also envision a scenario in which an intact capsid is docked at the dilated NPC, during which external perturbations constrict the central channel. In this case, the docked capsid will encounter additional inward stress, which may rupture the capsid lattice. Multiple conformational states of constituent NPC subcomplexes can be modeled with a multiconfiguration coarse-graining (MCCG) framework, in which the degree of coupling between different CG conformational states reproduces the underlying all-atom free energy surface projected onto the CG coordinates (66). It will be valuable to incorporate these sophisticated features in our CG NPC model based on experimental structural data to investigate how dynamic variation of NPC conformation modulates dynamics of translocation and capsid structural integrity.

To summarize, in this work, we elucidated key factors that can regulate the successful translocation of the HIV-1 viral capsid into the NPC central channel. We find that capsid morphology and orientation of approach are key factors in regulating a successful translocation event into the NPC central channel. A successful translocation event is contingent on several sequential steps. First, the tip of the capsid, when approaching from the cytoplasmic end, must establish significant contact with the central channel FG-NUPs. Second, the capsid translocates if the morphology is compatible with the dimension of the NPC central channel driven by the energetics of the interaction between central channel FG-NUPs and capsomers. Specifically, for the cone-shaped capsid, there is a gradient of the number of capsomers available to bind to the FG peptides of the NPC central channel due to the non-uniform cross-section. However, the gradually increasing cross-section also presents an incremental steric barrier impeding translocation. We find that the progressive relaxation of the inner ring scaffold leading to the dilation of the channel serves to relieve the resulting steric barrier for translocation (49). Finally, we found that extended (not associated) configurations of disordered NUP98 as observed *in vivo* when tethered to the NPC promote capsid association to the NPC central channel. Our results indicate that energetically favorable interactions between FG-NUP98 and capsid reduce the initial association barrier of the capsid to the NPC central channel.

Structural analysis of the viral capsids associated to the NPC central channel also indicates the appearance of highly correlated striated patterns of lattice disorder. These lattice disorder patterns are indicative of both a capsid lattice plasticity response and the weakening of the capsid structural integrity as a consequence of the radial stress from the spatial confinement of the NPC central channel and internal RNP complex. Once the capsid tip reaches the nuclear end of the NPC, the binding of the host factors NUP153 and CPSF6 mediate the nuclear import (25, 67). Furthermore, NUP153 and CPSF6 form higher-order aggregates templated by the viral capsid (68, 69). How these host factors regulate the dynamics of capsid passage from the NPC central channel to the interior of the nucleus and modulate the capsid structural integrity will be the subject of future investigation.

## METHODS

### CG model of the NPC

We first determined the CG mapping of NUP monomers from all-atom (AA) MD simulation trajectories using the Essential Dynamics Coarse-Graining (EDCG) method (35) to identify the group of consecutive residues that maximized the collective motions of the mapped CG protein with the motions of the corresponding AA trajectory. All NUP monomers were mapped such that the average AA to CG mapping resolution is ∼ 5 amino acid residues per CG site. We then determined the intramonomer bonding topology from the same AA MD trajectory by creating a heterogeneous elastic network model (hENM) (36).To create the intramonomer bonding topology, we considered all CG sites within 3 nm of a central CG site. To evaluate whether the CG hENM models recapitulate the AA fluctuations, we compared the root mean square (RMS) fluctuations of each CG NUP monomer with the reference AA MD simulations. To model NUP subcomplexes (Y-complex dimer, NUP54-NUP58-NUP62, and NUP155-NUP93) we created a weak harmonic bonding network with a force constant of 0.01 kcal mol^-1^Å^-2^ between neighboring monomers with a distance cutoff of 3 nm. The intermonomer force constant value was phenomenologically chosen to preserve the shape of each subcomplex.

Intermolecular non-bonded CG interactions between NUP monomers were modeled using a combination of repulsive excluded volume and attractive interactions. The repulsive excluded volume interaction (*E*_*excl*_) was modeled with a soft cosine potential,

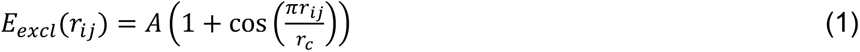

where *r*_*ij*_ is the pairwise distance between CG site types *i* and *j*. The value of *A* was set to 15 kcal/mol for all *ij* pairs. The distance cutoff (*r*_*c*_) for the excluded volume interactions was set at 1.0 nm. Attractive interactions between NUP subcomplexes at the key binding interactions were modeled with pairwise Gaussian potential (*E*_*gauss*_),

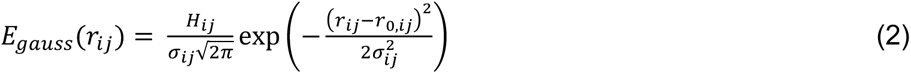

where *r*_0,*ij*_ and *σ*_*ij*_ are the mean and standard deviation of the distance between CG site types *i* and *j*. First, we determined the CG site types *i* and *j* that are in close contact in the mapped AAMD trajectories of NUP heterodimer complexes with a distance cutoff of 1.75 nm and standard deviation less than or equal to 0.12 nm. For all the subset of CG *ij* pairs in close contact, *r*_0,*ij*_ is determined from the mapped AA MD trajectories from the first peak of the pair correlation functions. For all CG *ij* pairs a *σ*_*ij*_ value of 0.12 nm was used. The constant *H*_*ij*_ for each binding interface was optimized using Relative Entropy Minimization from the atomistic trajectories of the respective all-atom subcomplexes (37). The derived attractive interactions (*E*_*attr*_) were validated by comparing the pair correlation functions between selected CG (*ij*) pairs of the CG NPC model and AA simulations of the corresponding heterodimer complexes at the binding interfaces.

Membrane association interactions between the CG sites of the NPC and CG lipid head group were modeled using a 12-6 Lennard-Jones potential (*E*_*scLJ*_) with a modified soft-core (70),

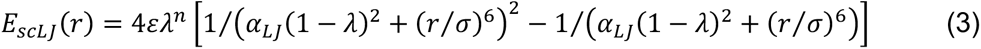

where, *n* = 2, *α_LJ_* = 0.5, *α* = 0.6, and *σ* = 1.5 nm. A minimum protein-lipid interaction strength (*ε* = 1.5 kcal mol^-1^) was identified and used in our simulations. Specifically, protein-lipid interactions were added to CG sites of the *β*-propeller domain of NUP155, NUP133, and NUP160.

The CG NUP98_1-620_ chains consist of 124 beads (**Fig. S15**). The CG model of the folded GLEBS domain (residue 157-213) was derived from AA MD simulation trajectory using EDCG and hENM protocols. The initial atomistic configuration of the GLEBS domain was generated using Alphafold2 (71). The rest of the CG NUP98 (residue 1-156 and residue 214-620) was represented as a linear polymer chain to mimic the disordered conformation. Each CG bead of the polymer chain is linked with a flexible harmonic with equilibrium bond length of 2 nm and harmonic force constant of 0.5 kcal/mol Å^−2^. The CG beads of the FG-rich region (residue: 1-156 and residue: 214-480) interact within the same chain and with other chains through 12-6 Lennard-Jones potential (*E*_*scLJ*_) with a modified soft-core. Here, *n* = 2, *α*_*LJ*_ = 0.5, *α* = 0.6, and *σ* = 1.25 nm are the parameters used. The strength of the inter- and intra-chain interactions (*ε*_*NUP*98_) between the FG-rich fragments of NUP98 is varied from 0.3 to 1.5 kcal/mol. The interaction between FG-motif sites of NUP98 and CA monomers is the same as between FG-motif sites of NUP62 and CA and is modeled with attractive pairwise Gaussian interactions (*E*_*gauss*_). All other intermolecular non-bonded interactions between NUP98 and other components are modeled with repulsive excluded volume interactions.

### CG model of HIV-1 Capsid

We derived a “bottom-up” CG model of CA from AA MD trajectories of a composite system of three CA hexamers complexed with IP6 and FG peptide. The CG CA monomer model was derived from the AA MD trajectory using the EDCG method (35). The CA monomer contains 47 CG sites with a resolution of ∼ 5 AA residues per CG site. The number of CG sites was chosen to balance the computational efficiency of simulating a full capsid and least-squared error in the principal component subspace. We mapped each FG peptide into a single CG site by calculating the center of mass of the 12-residue peptide bound to the CA monomer. The initial configuration of the FG peptide in the composite AA complex was generated from PDB 5TSX. After CG mapping of each component, the intramonomer bonding topology of the CA monomer was generated using hENM with a cutoff of 3 nm (36).

Intermolecular non-bonded CG interactions between CA monomers were modeled using a combination of repulsive excluded volume (*E*_*excl*_) and attractive pairwise Gaussian interactions (*E*_*gauss*_). The value of *A* in *E*_*excl*_ was 15 kcal/mol for all *ij* pairs of CA. The distance cutoff (*r*_*c*_) for the excluded volume interactions was set at 1.0 nm. To derive the attractive interactions (*E*_*gauss*_) we assumed that all CG beads of CA monomers that are in close contact participate in associative interactions. We classified close contact between CA CG sites in the CG mapped AA MD trajectory of the three CA hexamer complex as the *ij* pairs within a distance cutoff of 1.75 nm and a standard deviation of less than 0.15 nm. The parameters *r*_0,*ij*_ and *σ*_*ij*_ were determined by a fit to the first peak of the pair correlation function between CG *ij* pairs through least-squares regression. The constant *H*_*ij*_ for CG *ij* pairs was optimized using Relative Entropy Minimization from the corresponding AA MD trajectory (37). The derived CG parameters were evaluated by simulating full capsids of different shapes (conical, pill, and ellipsoid) and monitoring whether the capsid lattice order was maintained during the timescales of the simulations. Attractive interaction between the CG FG site and CA monomer was derived using the identical protocol described earlier. For all CG *ij* pairs for the FG-CA attractive interactions (*E*_*gauss*_) the *σ*_*ij*_ value of 0.12 nm was used.

### CG model of Ribonucleoprotein Complex (RNP)

The CG RNP in our simulations was modeled as a linear polymer chain containing 3000 beads. Each RNP polymer in our simulation minimally represents a 9-kb RNA genome complexed with nucleocapsid proteins. The mass of each CG site in the RNP was set to 500 Da. Each CG bead of the RNP was linked with flexible harmonic bonds with a harmonic force constant of 0.5 kcal/mol Å^−2^. The CG beads of the RNP interact with each other through a 12-6 Lennard-Jones potential (*E*_*scLJ*_) with a modified soft-core. Here, *n* = 2, *α*_*LJ*_= 0.5, *α* = 0.6, and *σ* = 1.25 nm are the parameters used. The strength of the RNP-RNP interaction (*ε*) was varied from 0.3 to 0.9 kcal/mol, which allows for modulating the condensation state of the RNP complex as noted earlier. Intermolecular non-bonded CG interactions between CA and RNP polymer were modeled with a combination of the repulsive excluded volume (*E*_*excl*_) and attractive pairwise Gaussian interactions (*E*_*gauss*_). The parameters for the repulsive excluded volume interactions are identical to that between CA and CA. The RNP polymer interacts with two C-terminal CG beads of the CA through attractive pairwise Gaussian interactions (*E*_*gauss*_) to emulate electrostatic interactions between the RNP and charged residues in the C-terminal end of CA (*H*_*ij*_ = - 4.5 kcal/mol, *r*_0,*ij*_ = 1.25 nm, and *σ*_*ij*_ = 0.12 nm).

### CG model of Lipids

A 4-site CG lipid model was used consisting of 1 head, 1 interfacial, and 2 hydrophobic tail beads using the same functional form as in ref. (38). We note that an identical lipid bilayer model was used to simulate other complex biological systems, such as immature virion assembly and SARS-CoV-2 virion (31, 34, 72).

### CG simulations

All CG MD simulations of composite NPC, lipid, and capsid were prepared and simulated in the LAMMPS MD software(73). Two preequilibrated CG lipid bilayers were first aligned coplanar (in the *x* and *y* direction) to the cytoplasmic and nuclear rings of the CG NPC model. All CG lipids within 0.5 nm of the cytoplasmic and nuclear rings were deleted to avoid unphysical contact. Similarly, the CG lipids located in the NPC channel (coplanar to CR and IR) were also deleted. A cylindrical membrane pore was also created, which wraps around the NPC IR. The composite membrane system was first minimized and then equilibrated for 10 × 10^6^ *τ*_*CG*_ by integrating only the CG lipids (the position of the NPC CG beads was constrained to the initial position) to allow relaxation. The system was then further equilibrated by 10 × 10^6^ *τ*_*CG*_. In these simulations, the membrane binding *β*-propeller domain of NUP155, NUP133, and NUP160 were also integrated along with the lipids, while the position of the remainder of the NPC was kept constrained. In 20 × 10^6^ *τ*_*CG*_ of equilibration, all *β*-propeller domains of NUP155, NUP133, and NUP160 were associated with the lipid. The final dimension of the simulation cell was 200 nm in both the *x* and *y* direction. In the *z* direction, the dimension of the simulation cell was 300 nm. All CG MD simulations were performed using a CG MD time step (*τ*_*CG*_) of 50 fs, and the equations of motions were integrated with the Velocity Verlet algorithm. All simulations were periodic in all three directions. Final production simulations were performed in the constant *Np*_*xy*_*T* ensemble. The temperature of the system was maintained using Langevin thermostat at 300 K with a coupling constant of 2000 *τ*_*CG*_ (74). The pressure of the system was maintained using a Nose-Hoover barostat with a coupling constant of 4000 *τ*_*CG*_ (75). Simulation trajectory snapshots were saved every 0.5 × 10^6^ CG MD timesteps.

Capsid translocation simulations were performed with a path-sampling approach as noted earlier. The simulations were discretized into consecutive (*n*) segments. For each segment, 5 replica simulations, each 40 × 10^6^ CG timesteps long, were performed. For each segment, the probability distribution of *D*_*Cap-NPC*_ calculated from final 5 × 10^6^ CG timesteps for all 5 replica simulations were fitted to a normal distribution. The endpoint of a trajectory that yielded the *D*_*Cap-NPC*_ value closest to the mean of the cumulative probability distribution was used to initialize the next simulation segment. Finally, the selected trajectory from each segment was concatenated to create the complete trajectories of capsid translocation. Simulations of freely diffusing capsids were performed by initially placing a capsid in a simulation cell of dimension 200 nm in *x*, *y*, and *z* directions. The initial position of the capsid was adjusted such that the geometric center of the capsid was aligned with the center of the simulation cell. These simulations were performed in the constant *NVT* ensemble. The temperature of the system was maintained using a Langevin thermostat at 300 K with a coupling constant of 2000 *τ*_*CG*_.

### CG Model Analysis

Visualization of the simulation trajectories was performed in VMD (76).

a. *Capsid-FG association*: We identify CA-FG association by using the following distance-based criteria:

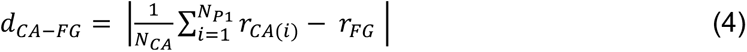 where *d*_*CA-FG*_ is the distance between the selected FG site and center of mass of the CA monomer calculated over selected indices. Here, *r* is the coordinate of the CG sites and *N*_*CA*_ is the total number of CG sites considered for the distance calculation. For the calculation, CA monomer CG site indices 9-14, 21-23, and 26-28 are considered. Note that a CA monomer consists of 46 CG sites. A CA monomer and FG site of NUP62 is associated if *d*_*CA-FG*_ < 3 nm.
b. *Capsid lattice order analysis*: We quantified the capsid lattice order by the neighbor averaged Steinhardt bond order parameter (41, 42). The Steinhardt bond order parameters determine the order of the local environment using an algorithm based on spherical harmonics. To calculate the averaged Steinhardt bond order parameter, we considered the CG site index 21, which is the closest site to the geometric center of the CA monomer. Considering the selected CG site for each CA monomer of the viral capsid, the neighbor averaged Steinhardt bond order parameter

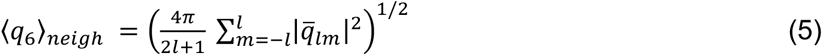 where 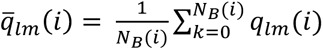 and *l* = 6. Here *N*_*B*_ is the number of neighbors of CA monomer *i*, and the reference CA monomer itself. The term 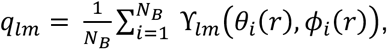 and ϒ_*lm*_(*θ_i_*(*r*), *ϕ_i_*(*r*)) are spherical harmonics of rank *l* and *m*, where *θ*_*i*_(*r*) and *ϕ_i_*(*r*) are the polar angles of each of the *N*_*B*_ bonds between the central CA monomer and neighboring CA monomers. We note that the closest 3 neighbors of CA monomer *i* are considered for the calculation of 〈*q*_6_〉_*neigh*_.
c. *Largest disordered cluster calculation*: To investigate the formation of disordered domains of capsid lattice during translocation through the NPC central channel, we performed clustering analysis. First, we identified all CA monomers with 〈*q*_6_〉_*neigh*_ < 0.4. Then for a particular CA monomer with 〈*q*_6_〉_*neigh*_ < 0.4 we identified any closest 3 neighbors with 〈*q*_6_〉_*neigh*_ < 0.4. The largest disordered cluster was defined as the cluster containing the most contiguous CA monomers with 〈*q*_6_〉_*neigh*_ < 0.4 for each configuration. The radius of gyration (*R*_*g*_) of the largest cluster was calculated for the largest connected disordered CA domain.

## DATA AVAILABILITY

The CG model parameters of the NPC and HIV-1 capsid, and input files are provided at the https://doi.org/10.5281/zenodo.8217689. Additional details of the CG model are provided within the article or Supporting Information.

## ACKNOWLEDGEMENTS

The computational resources were provided by the Texas Advanced Computing Center (TACC) at The University of Texas at Austin and the Research Computing Center (RCC) at The University of Chicago. Simulations were performed using resources provided by the Extreme Science and Engineering Discovery Environment (XSEDE) (prior to September 2022) (77), supported by the National Science Foundation grant number 1548562, by the Advanced Cyberinfrastructure Coordination Ecosystem: Services & Support (ACCESS) program (after September 2022), which is supported by National Science Foundation grants numbers 2138259, 2138286, 2138307, 2137603, and 2138296, and Frontera (at TACC) funded by the NSF (OAC-1818253).

## AUTHOR CONTRIBUTIONS

A. H. and G.A.V. designed the research. A. H. built systems, performed simulations, analyzed data, created figures, and prepared the initial manuscript draft. A.H. and G.A.V. contributed to the writing and editing of the final manuscript.

Conceptualization: A. H., G. A. V.
Methodology: A. H.
Investigation: A. H.
Resources: A. H., G. A. V
Visualization: A. H.
Supervision: G. A. V.
Writing—original draft: A. H.
Writing—review & editing: A. H., G. A. V.
Funding acquisition: G. A. V.

