## Supplementary Material for "HIV-1 Capsid Shape, Orientation, and Entropic Elasticity Regulate Translocation into the Nuclear Pore Complex"


### Supporting Methods

#### S1. Deriving coarse-grained (CG) model and molecular interactions from atomistic simulations using "bottom-up" methods.

We systematically derived CG models and interactions of NUP monomers and subcomplexes from reference all-atom MD simulations following a three-step hierarchical methodology. First, CG beads of NUP monomers were mapped from the corresponding all-atom simulation trajectories using the Essential Dynamics Coarse-Graining (EDCG) method (1). Using the derived CG map, we represented intra-protein interactions using heterogeneous elastic network models (hENMs) (2). Then attractive short-ranged CG inter-protein interactions were derived from all-atom simulation trajectories of respective NUP subcomplexes using relative entropy minimization (3). These methods have been successfully utilized before to derive "bottom-up" CG models of various multicomponent protein assemblies (4-6). The details of the coarse-grained modeling procedures are described below.

#### 1. CG mapping from reference atomistic simulations.

To define the CG sites of NUP monomers from atomistic simulation trajectories, we used Essential Dynamics coarse-graining (EDCG) (1). EDCG is a "bottom-up" coarse-graining procedure in which $C_\alpha$ sites of a number of consecutive all-atom residues are grouped to a CG site such that the principal modes of motion sampled during atomistic simulations are conserved. In EDCG, the mapping operator ($\boldsymbol{M}_{\mathbf{R}}^{N}\colon \mathbf{r}^{n} \rightarrow \mathbf{R}^{N}$), is variationally optimized using simulated annealing to obtain the global minimum of the target residual ($\chi^2$). Here, $\mathbf{r}^n$ and $\mathbf{R}^N$ are the configurations of the atomistic and coarse-grained trajectories, respectively. The all-atom to coarse-grained mapping function is adjusted during the optimization to minimize the target residual ($\chi^2$):

$$\chi^2 = \frac{1}{3N}\sum_{I=1}^{N} \langle \sum_{i,j} \left|\boldsymbol{r}_i - \boldsymbol{r}_j\right|^2 \rangle_t \quad : \; i,j \in I, j \geq i \tag{1}$$

where, $N$ is the total number of CG sites. Here $i,j$ are the unique pairs in the group of all-atom residues that are part of the CG site, $I$. $\boldsymbol{r}_i = \boldsymbol{x_i} - \langle \boldsymbol{x_i} \rangle_t$ is the displacement of atom $i$ from the atom's mean position, $\langle \boldsymbol{x_i} \rangle_t$. When atoms $i,j$ move in correlated fashion in the atomistic trajectory, then $\left|r_i - r_j\right|^2$ is small. For $N$ CG beads ($N$ all-atom residue segments), $N-1$ segment boundaries are initially defined along the primary amino acid sequence. At each step of simulated annealing, a new CG map is defined by moving the position of the segment boundary atoms along the primary sequence. The new map is either accepted or rejected according to a Metropolis-Hastings criterion. The new map is accepted if the value of the residual is less than its predecessor

($\chi_1^2 < \chi_0^2$). If the new CG map has a higher residual, then the map is accepted with a probability $exp(-\Delta\chi^2/T)$. Here $T$ is the coupling to a temperature parameter that is initially high to allow the boundary atoms to move randomly and allow escape from local minima and is gradually lowered during the optimization. All CG maps of NUP monomers were generated from the final 500 ns of atomistic MD trajectories and had an average resolution of ~ 5 $C_\alpha$ residues per CG site.

**2. Intra-protein bonded interactions derived from atomistic MD simulations.**

Intra-protein bonded interactions were defined using heterogeneous elastic network model (hENM), which captures the flexibility of the protein (2). In the hENM method, initially, harmonic bonds with identical force constant ($k_{ij}$) are assigned between the central CG site ($i$) and all the neighboring CG sites ($j$) within a user-specified distance cutoff ($r_{cut}$). The harmonic force constant ($k_{ij}$) for each bond is iteratively optimized until the fluctuations in the CG model converge to that of reference atomistic simulation trajectory, i.e.,

$$\frac{1}{k_{ij}^{n+1}} = \frac{1}{k_{ij}^{n}} - \alpha\left(\langle r_{ij}^2\rangle_{\mathrm{CG}} - \langle r_{ij}^2\rangle_{\mathrm{AA}}\right) \tag{2}$$

where $k_{ij}^n$ and $k_{ij}^{n+1}$ is the harmonic force constant at the iteration step $n$ and $n+1$ step, respectively. $\langle r_{ij}^2\rangle = \langle\left(x_{ij} - \langle x_{ij}\rangle\right)^2\rangle$ is the mean-squared fluctuation for each CG site pair $i,j$. $\alpha$ is a parameter that controls the magnitude of the adjustment for each iteration. For all NUP monomer CG models, we used a cutoff distance ($r_{cut}$) of 3 nm. The optimization procedure to derive the force constant ($k_{ij}$) was performed from the final 200 ns of atomistic MD trajectories.

**3. Interprotein short-ranged attractive interactions from reference atomistic simulations.**

Inter-protein attractive interactions between CG sites of NUP subcomplexes are modeled with pairwise short-ranged Gaussian potentials ($E_{gauss}$),

$$E_{gauss}(r_{ij}) = \frac{H_{ij}}{\sigma_{ij}\sqrt{2\pi}} \exp\left(-\frac{\left(r_{ij}-r_{0,ij}\right)^2}{2\sigma_{ij}^2}\right) \tag{3}$$

where $r_{ij}$ and $\sigma_{ij}$ are the mean and standard deviation of a CG site pair $i,j$. The constant $A_{ij}$ is optimized using Relative Entropy Minimization (REM) (3). The constant $A_{ij}$ (referred to as $\lambda$ in equation 4) is iteratively optimized using Newton-Raphson method as follows,

$$\lambda_{n+1} = \lambda_n - \chi \frac{\left(\frac{\partial S}{\partial \lambda}\right)}{\left(\frac{\partial^2 S}{\partial \lambda}\right)} \tag{4}$$

$\chi$ is the "learning rate" of the iterative optimization procedure. Note that the "learning rate" ($\chi$) is varied throughout the optimization procedure.

To perform the REM iteration cycle, we first created a CG NUP subcomplex by mapping the final configuration from the all-atom trajectory. All the NUP subcomplexes used to derive inter-protein interactions consist of two NUP monomers that are in direct contact in the cryo-electron tomography (cryo-ET) structural data (7). For each iteration, a CG simulation of the NUP subcomplex was evolved for 25 × $10^6$ CG MD timesteps at 300 K. Total 500 step iteration cycle was performed as changes between successive steps are effectively zero within 500 steps. Learning rate ($\chi$) is varied as 0.5 (1-100 steps), 0.1 (101-200 steps), 0.01 (201-500 steps).

**S2. Atomistic simulation of NUP monomers and heterodimers.**

We performed 1000 ns long atomistic MD simulations of NUP monomers and heterodimers in solution to derive the CG molecular model and interactions. We solvated the initial protein structure (monomers and heterodimers) in a cubic box of TIP3P water. The solvated system was built such that there is at least 1.2 nm layer of water between the protein surface and the edge of the simulation cell. To achieve a physiological ion concentration of 150 mM, we added $Na^+$ and $Cl^-$ ions to the solvated system by replacing randomly selected TIP3P water molecules. The simulation system was then energy minimized using the steepest descent method until the target maximum force was less than 239 kcal/mol/nm. The system was equilibrated by applying harmonic positional restraints (spring constant value of 239 kcal/mol/nm$^2$) on protein heavy atoms for 500 ps in the constant NVT ensemble. The temperature of the system was maintained at 310 K using stochastic velocity rescaling thermostat with a time constant of 1 ps. A second constrained equilibration was done for 800 ps in the constant NPT ensemble at 310 K and 1 bar using the isotropic Parrinello-Rahman barostat with a 10 ps time constant. In this equilibration procedure, the harmonic positional constraint on protein heavy atoms was progressively decreased every 200 ps ($k$ = 239, 119.5, 23.9, 11.9 kcal/mol/nm$^2$). The production runs were performed in the constant $N_pT$ ensemble at 310 K and 1 bar. The temperature was maintained using Nose-Hoover chain thermostat with a 2 ps time constant. The pressure of the simulations was maintained with an isotropic Parrinello-Rahman barostat with a 10 ps time constant (8, 9). All simulations were performed with periodic boundary conditions in $x$, $y$, and $z$ directions. The protein was modeled with the CHARMM36m force field (10), and water was modeled with TIP3P parameters (11), using a 2 fs MD timestep. LINCS algorithm was used to constrain the bonds between heavy and hydrogen atoms (12). Electrostatic interactions were computed using the particle mesh Ewald method with a cutoff of 1 nm (13). The van der Waals force was truncated smoothly to zero between 1.0 and 1.2 nm. All atomistic simulations were performed using the Gromacs 2019 MD package (14).

### Supporting Figures

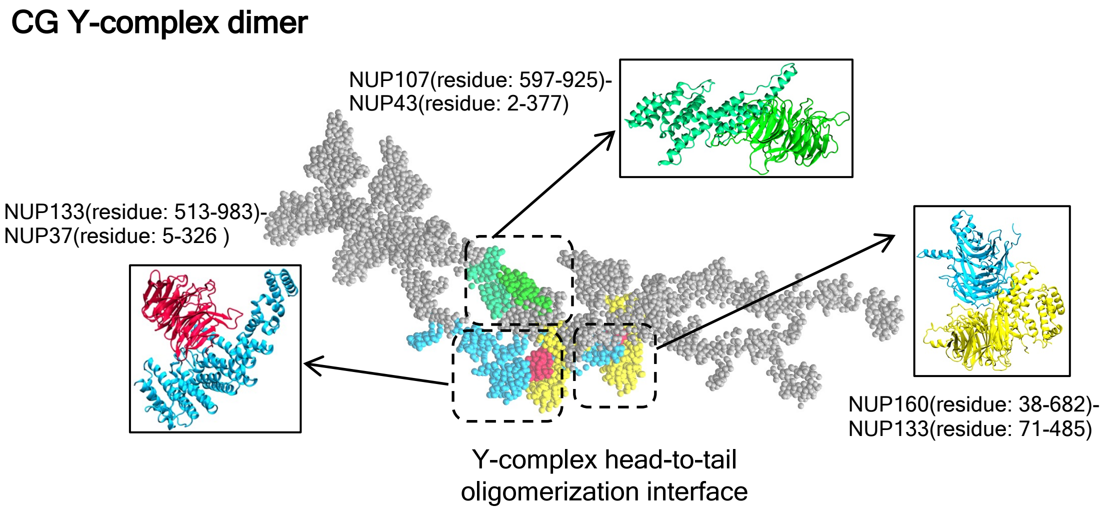


**Fig. S1. Representative depiction of the head-to-tail oligomerization interface between two adjacent Y-complex dimers.** The location of the binding interface between NUP160-NUP133, NUP133-NUP37, and NUP107-NUP43 complexes in the Y-complex dimer are labeled. The corresponding atomistic heterodimer complexes from which the CG molecular interactions are derived are shown in cartoon representation in the inset. Each atomistic heterodimer complex was evolved for 1000 ns to derive the corresponding CG molecular interactions. The NUPs at the binding interface are represented in the same color in Fig. 1 in the main manuscript. The rest of the CG beads of the Y-complex dimer is shown in gray spheres.

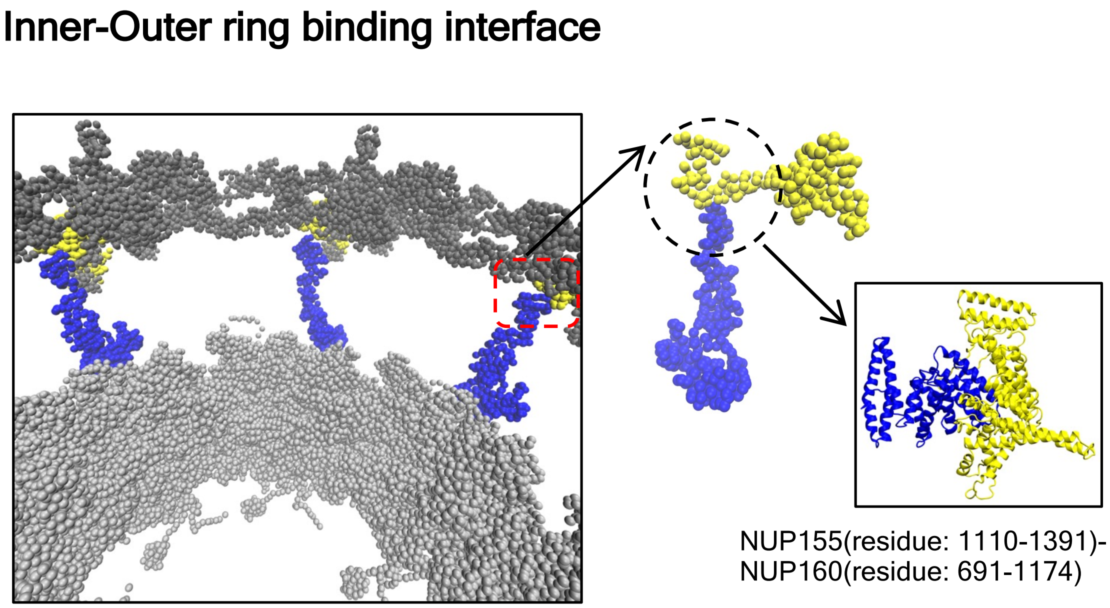


**Fig. S2. Representative depiction of the inner-outer ring binding interface.** 16 copies of NUP155 (blue spheres) are vertically oriented to bind NUP160 (yellow spheres). 8 copies of

NUP155 form a binding interface with NUP133 at the cytoplasmic and nuclear rings, respectively. The atomistic NUP155-NUP160 complex from which the CG non-bonded attractive interactions are derived is shown in cartoon representation in the inset. The atomistic NUP155-NUP133 heterodimer complex was simulated for 1000 ns to derive the corresponding CG molecular interactions. The NUPs at the binding interface are represented in the same color in Fig. 1 in the main manuscript. The rest of the CG beads are shown in gray spheres.

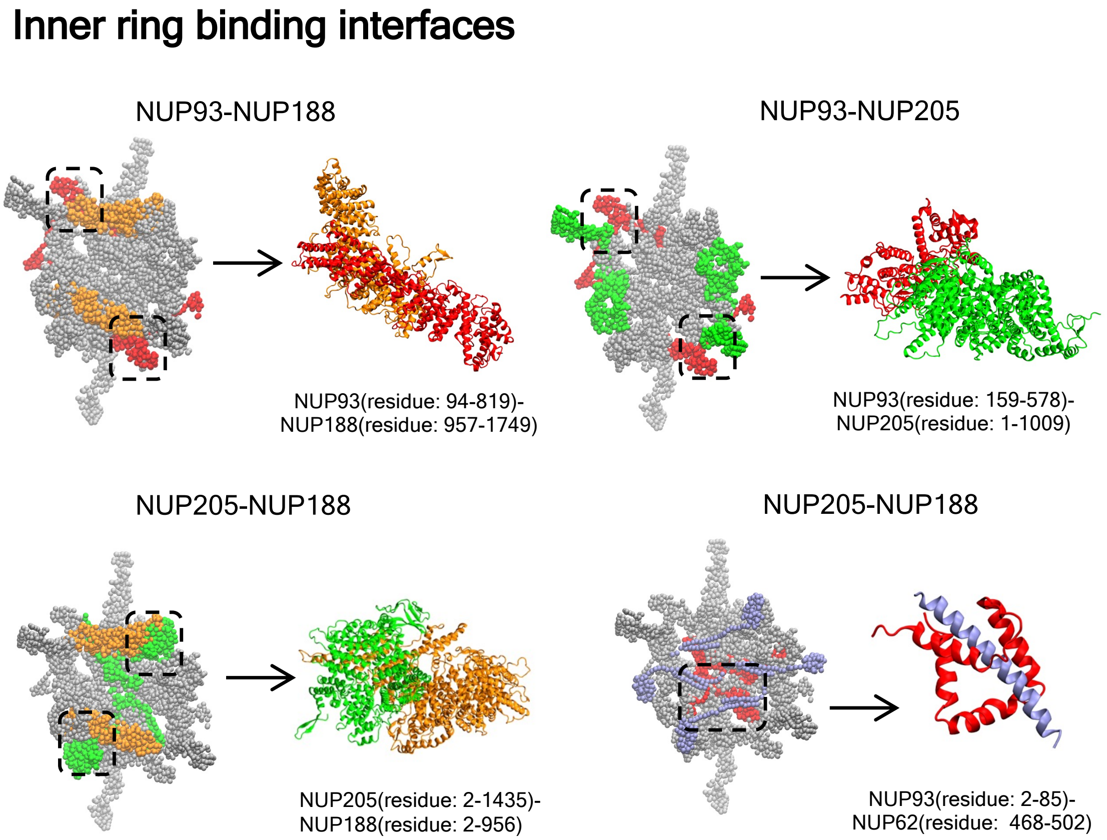


**Fig. S3. Representative depiction of the inner ring binding interfaces.** The location of the binding interface between NUP93-NUP188, NUP93-NUP205, NUP205-NUP188, and NUP93-NUP62 in an inner ring spoke is shown (left panel). The corresponding atomistic heterodimer complexes from which the CG molecular interactions are derived is shown in the right panel in the cartoon representation. Each atomistic heterodimer complex was simulated for 1000 ns to derive the corresponding CG molecular interactions. The NUPs at the binding interface are represented in the same color in Fig. 1 in the main manuscript. The rest of the CG beads of the inner ring spoke are shown in gray spheres.

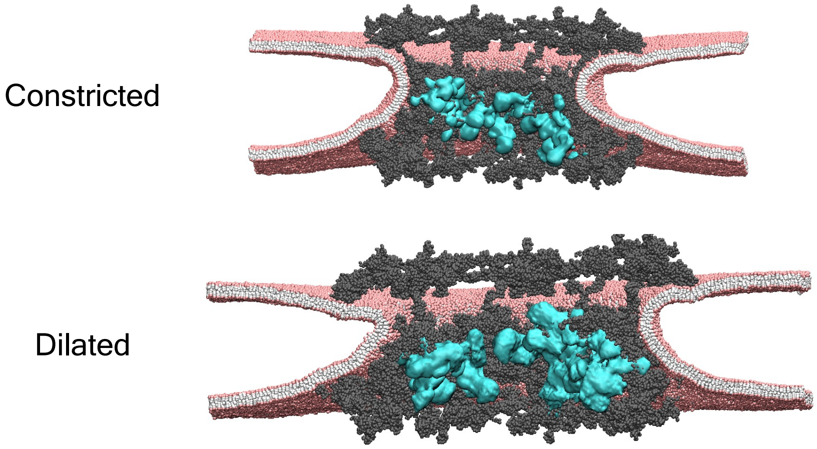


**Fig. S4. Time-averaged densities of the FG-NUPs of the NPC.** The density of the NUP54-58-62 heterotetramer complex were calculated from the final 150 × $10^6$ $\tau_{CG}$ of the CG MD simulations. The averaged density of the NUP54-58-62 heterotetramer complex is shown in cyan isosurface overlaid on the NPC. Rest of the NPC components are shown in gray spheres. The headgroup of the 4-site CG lipid is shown in pink spheres. The interfacial bead and two tail beads of the CG lipid are shown in white spheres.

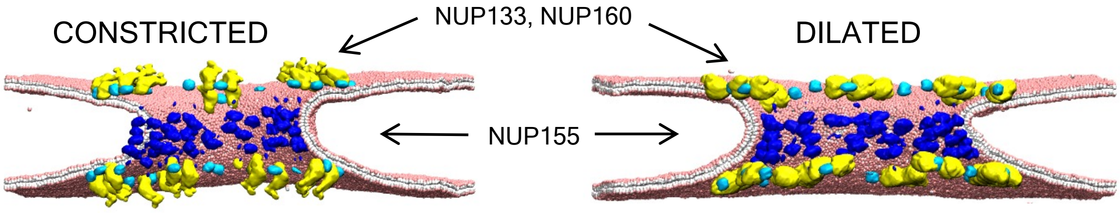


**Fig. S5. Time-averaged densities of the membrane-binding domains of the NPC.** The densities of each membrane-binding $\beta$-propeller domain are shown for the constricted and dilated NPC overlaid on the lipid bilayer. The densities were calculated from the final 150 × $10^6$ $\tau_{CG}$ of the CG MD simulations. The lipid bilayer is vertically sliced to show the density of NUP155 $\beta$-propeller domains (blue isosurface) bound to the membrane around the central channel. $\beta$-propeller domains of NUP133 and NUP160 are shown in cyan and yellow isosurface, respectively. The headgroup of the 4-site CG lipid is shown in pink spheres. The interfacial bead and two tail beads of the CG lipid are shown in white spheres.

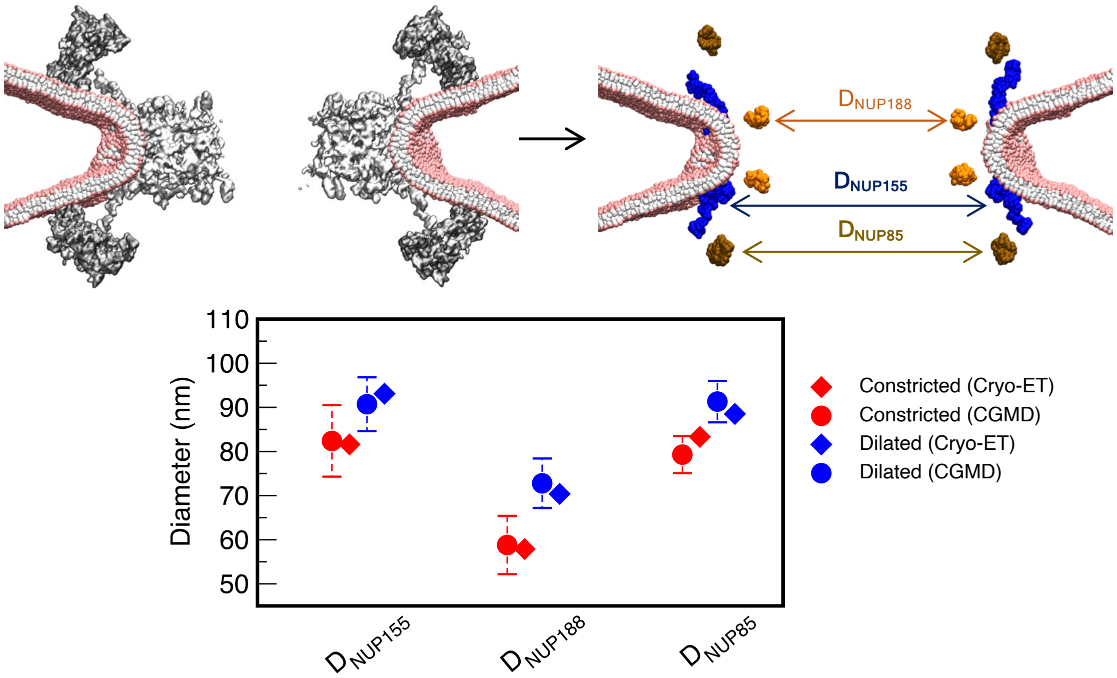


**Fig. S6. Analysis of NPC diameters measured between selected NUPs at the outer (cytoplasm and nuclear) and inner rings.** The outer ring diameters ($D_{NUP85}$) are estimated between the geometric center of NUP85 (inner tip of the Y-complex dimer). At the inner ring, diameters are measured between the geometric center of membrane-bound $\beta$-propeller domains of NUP155 ($D_{NUP155}$) and NUP188 ($D_{NUP188}$). The upper left panel shows the cutaway view of the NPC and lipid showing the outer (CR and NR) and inner rings. The upper left panel shows the same cutaway view highlighting only the NUP85, NUP155, and NUP188 proteins, between which the corresponding ring diameters are measured. The bottom panel shows the measured diameters for the constricted and dilated CG NPC models calculated from the final $150 \times 10^6$ $\tau_{CG}$ of the CG MD simulations. The same diameters measured for the reference cryo-ET structures are shown in the same plot.

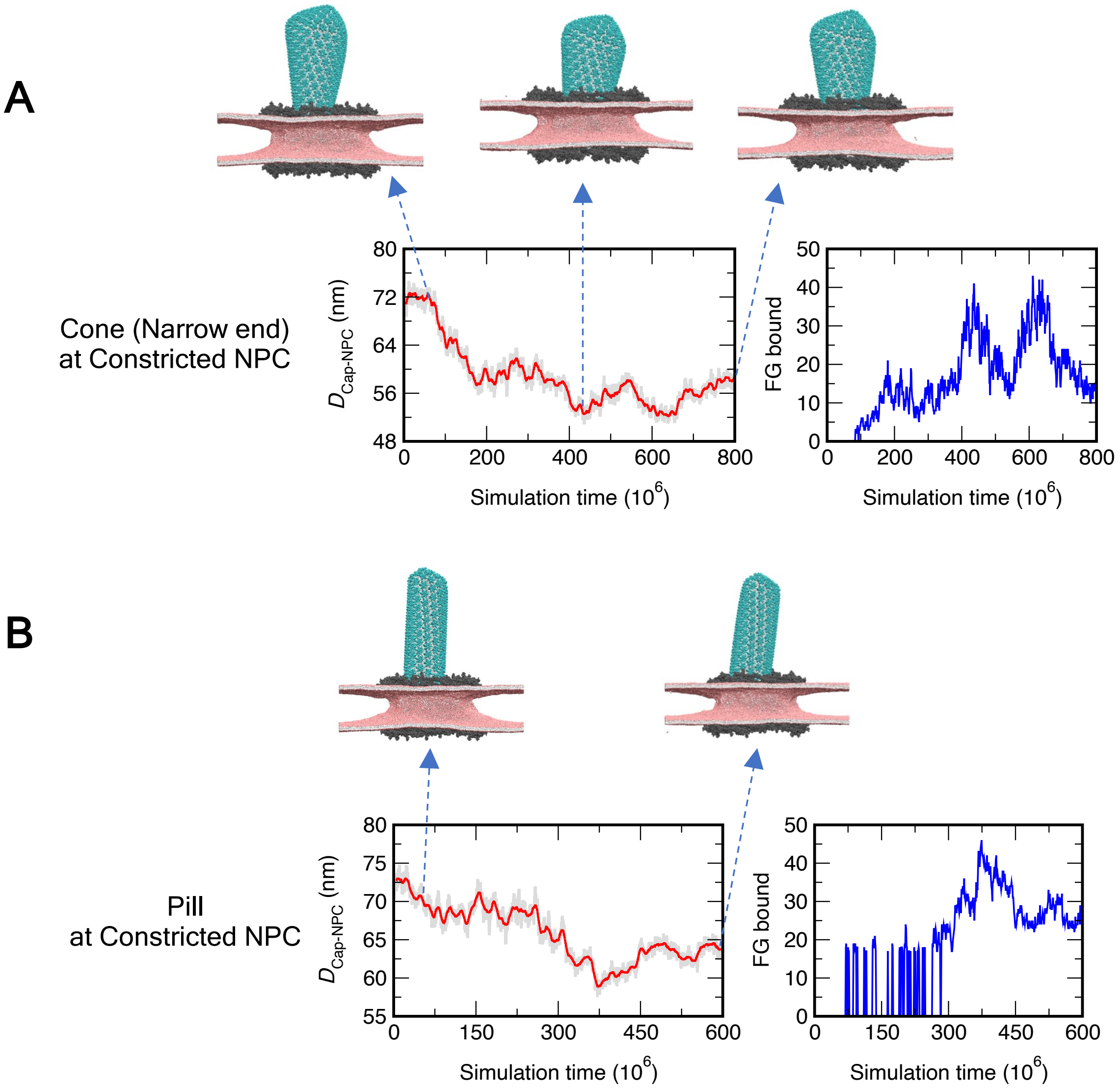


**Fig. S7. Translocation dynamics of HIV-1 capsid into the central channel of the constricted NPC.** Translocation dynamics time series plot (A and B) that depicts the distance ($D_{Cap-NPC}$) between the geometric center of the capsid and NPC inner ring (left panel in red), and the number of FG sites of NUP62 bound to CA (right panel in blue). The snapshots depict the cone-shaped (approaching from the narrow end) and pill-shaped capsid at the central channel of the constricted NPC at different points of the translocation dynamics trajectory. The capsid, NPC, and lipids are shown in the color scheme in **Fig. 3** of the main text.

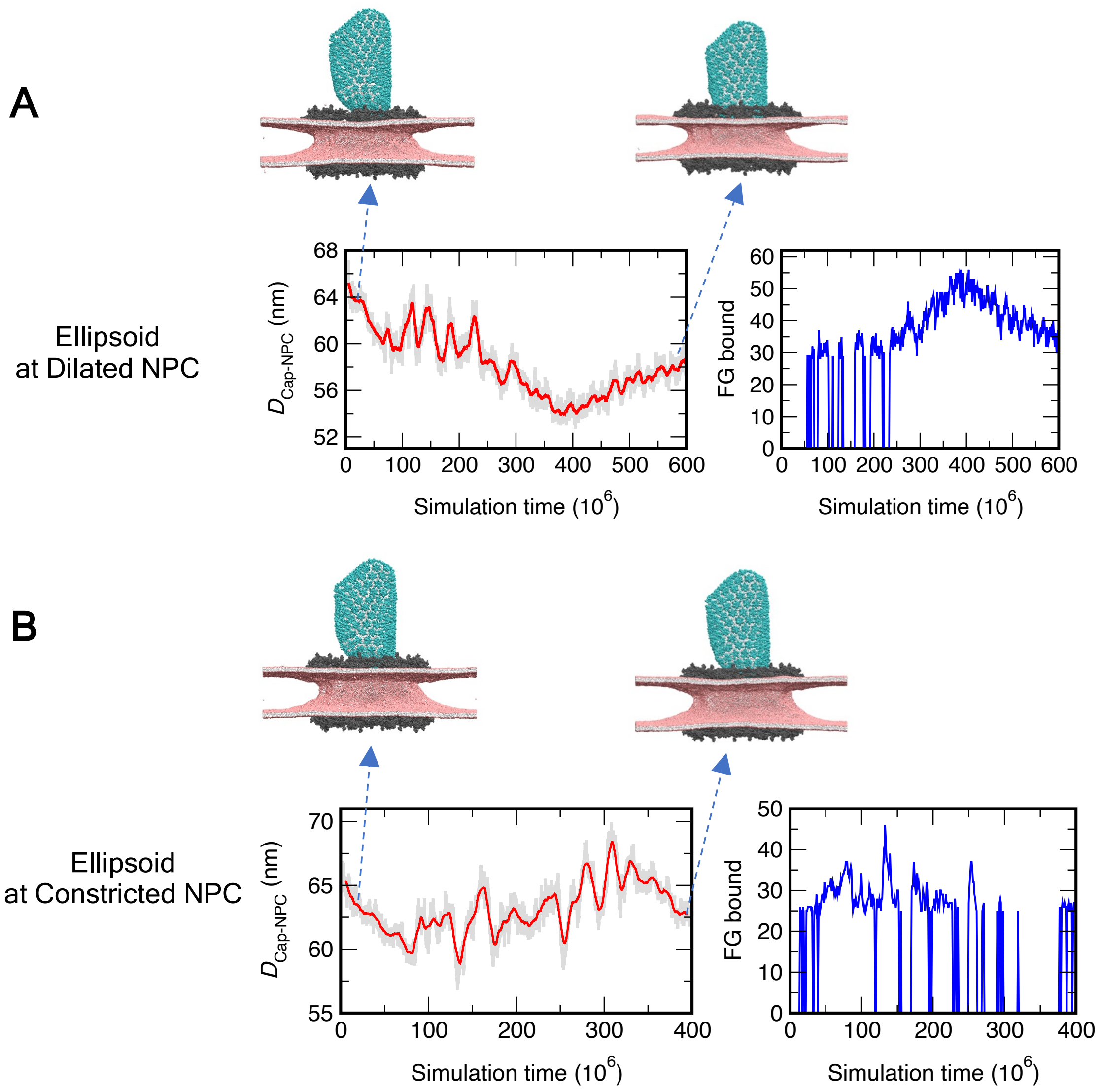


**Fig. S8. Translocation dynamics of ellipsoid-shaped capsid into the NPC central channel.** Translocation dynamics time series plot of the ellipsoid capsid into the dilated (A) and constricted (B) NPC. The capsid, NPC, and lipids are shown in the color scheme in **Fig. 3** of the main text.

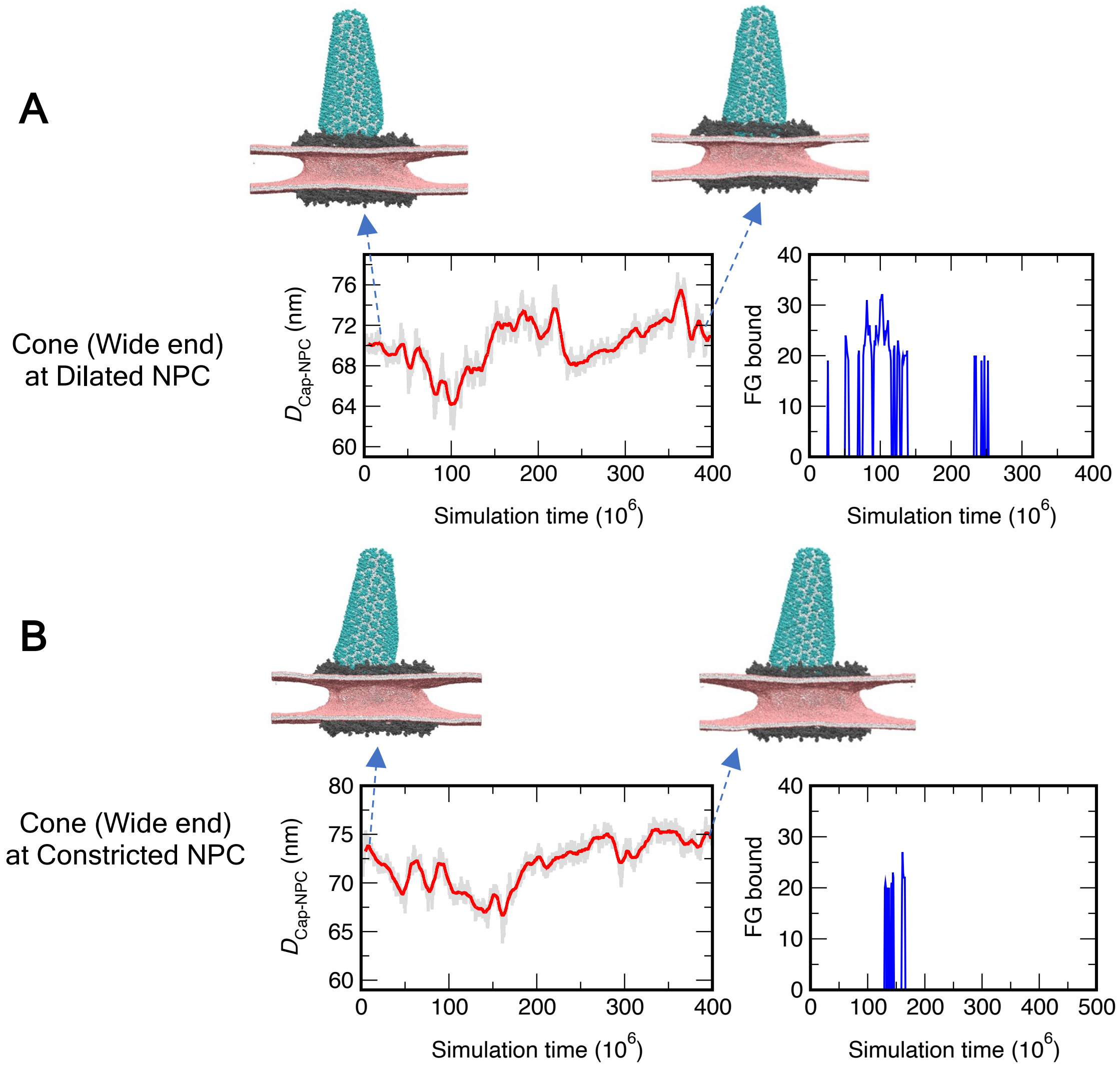


**Fig. S9. Translocation dynamics of cone-shaped capsid (approaching from the wide end) into the NPC central channel.** Translocation dynamics time series plot of the cone-shaped capsid into the dilated (A) and constricted (B) NPC. The capsid, NPC, and lipids are shown in the color scheme in **Fig. 3** of the main text.

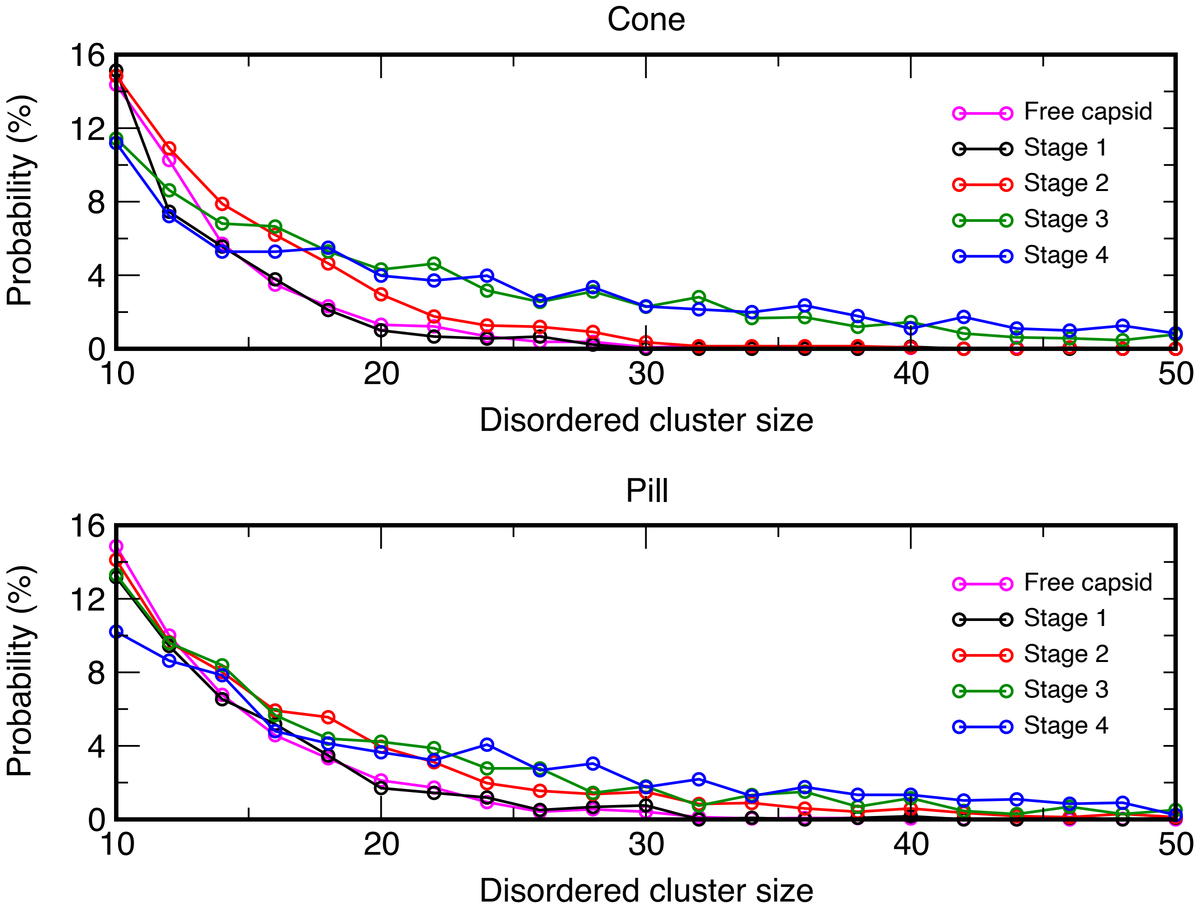


**Fig. S10. Distribution of the size of disordered CA cluster for the cone-shaped (*A*) and pill-shaped (*B*) at different stages of translocation into the dilated NPC central channel.** A particular CA monomer is classified as disordered if $\langle q_6 \rangle_{neigh} < 0.4$. Connected clusters of at least 10 CA monomers are considered.

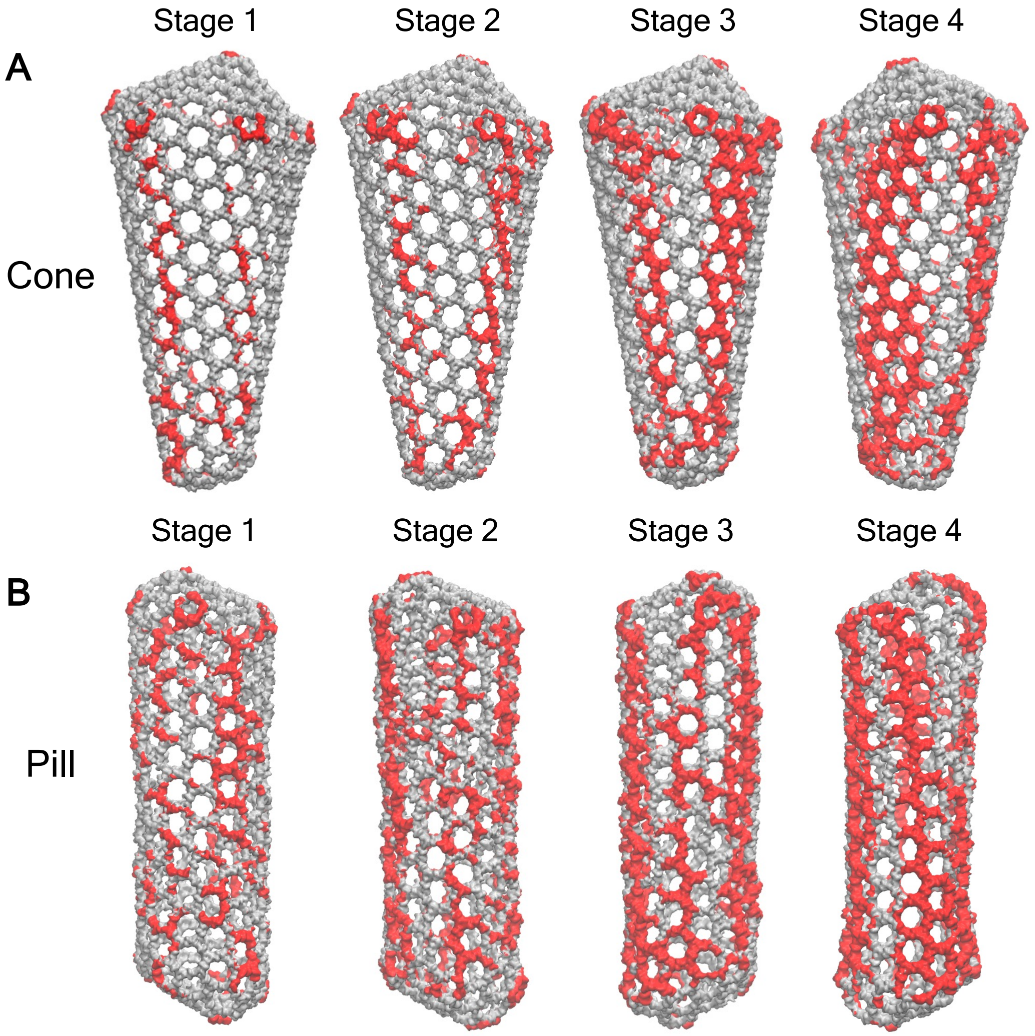


**Fig. S11. Time-averaged capsid lattice disorder at different stages of entry into the NPC central channel for the cone (*A*) and pill (*B*).** The CTD domain of capsid is shown in silver. The CTD domain of each CA monomer with $\langle q_6 \rangle_{neigh} < 0.4$ is shown in red. The CA NTD domain is not shown. We note that, the highlighted disordered domains (shown in red) are present in over 50% of the sampled trajectory frames.

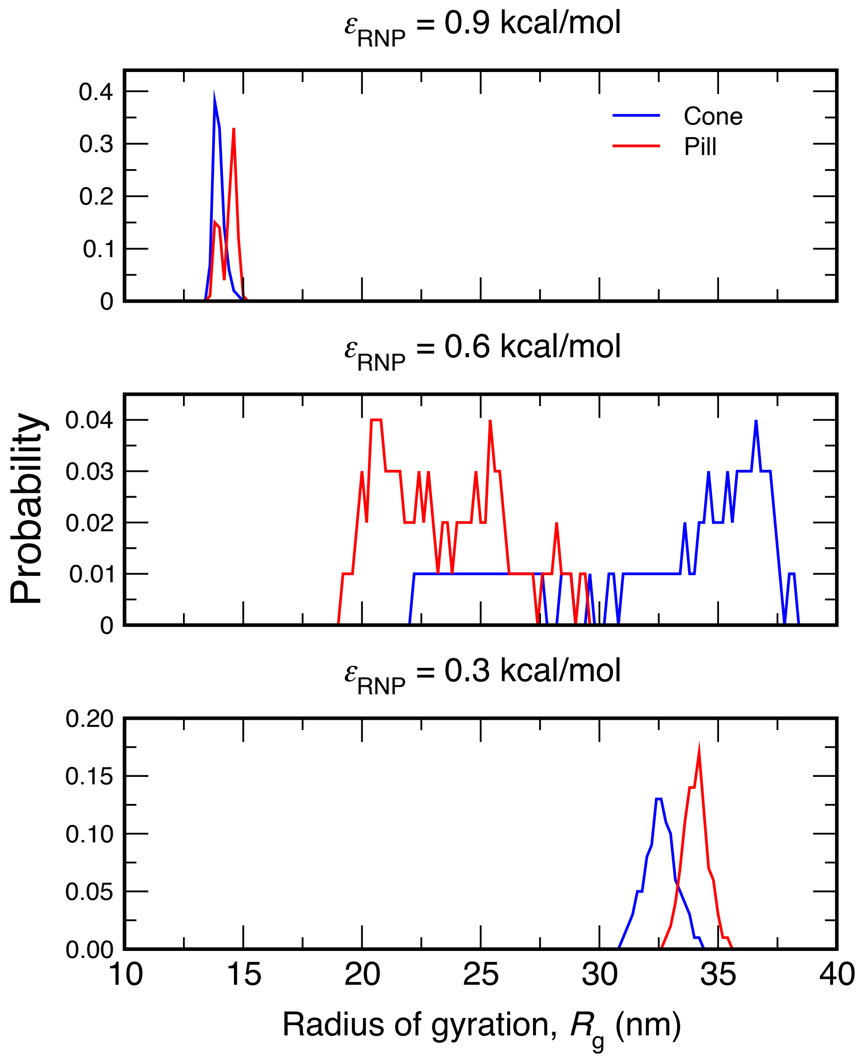


**Fig. S12. Probability distribution of the radius of gyration ($R_g$) of the RNP complex in the interior of the capsid.** From top to bottom. $\varepsilon_{RNP}$ is varied from 0.9 to 0.3 kcal/mol, modifying the RNP condensation states. For each $\varepsilon_{RNP}$, the $R_g$ distribution is shown in blue and red for the cone-shaped and pill-shaped capsid, respectively.

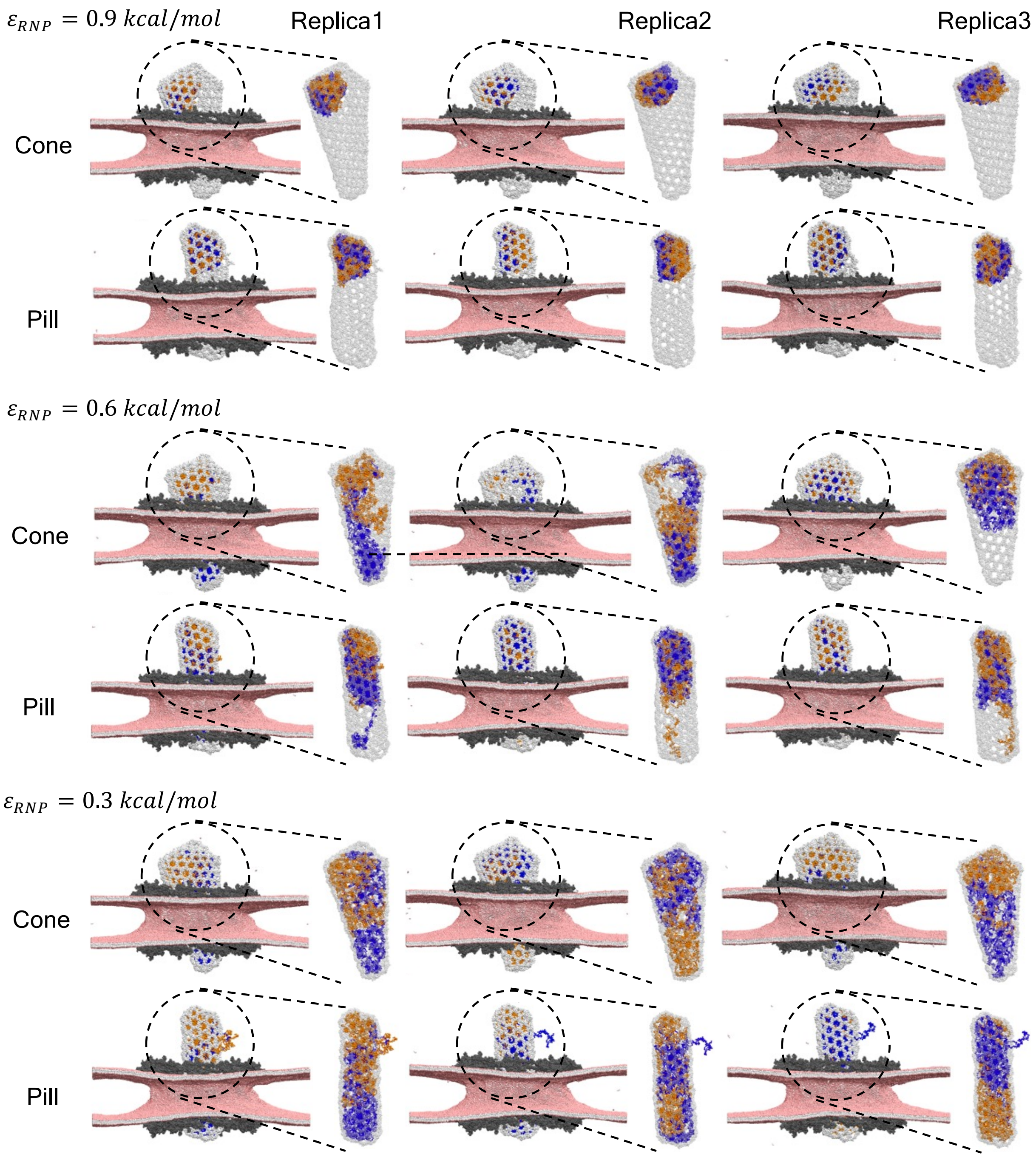


**Fig. S13. Snapshots of the RNP complex in the interior of the capsid.** The two RNP chains are shown in blue and orange spheres. The three panels next to each $\varepsilon_{RNP}$ correspond to 3 replica simulations. The capsid, NPC, and lipids are shown in the color scheme as in **Fig. 3**. The NTD domain of the capsid is not shown.

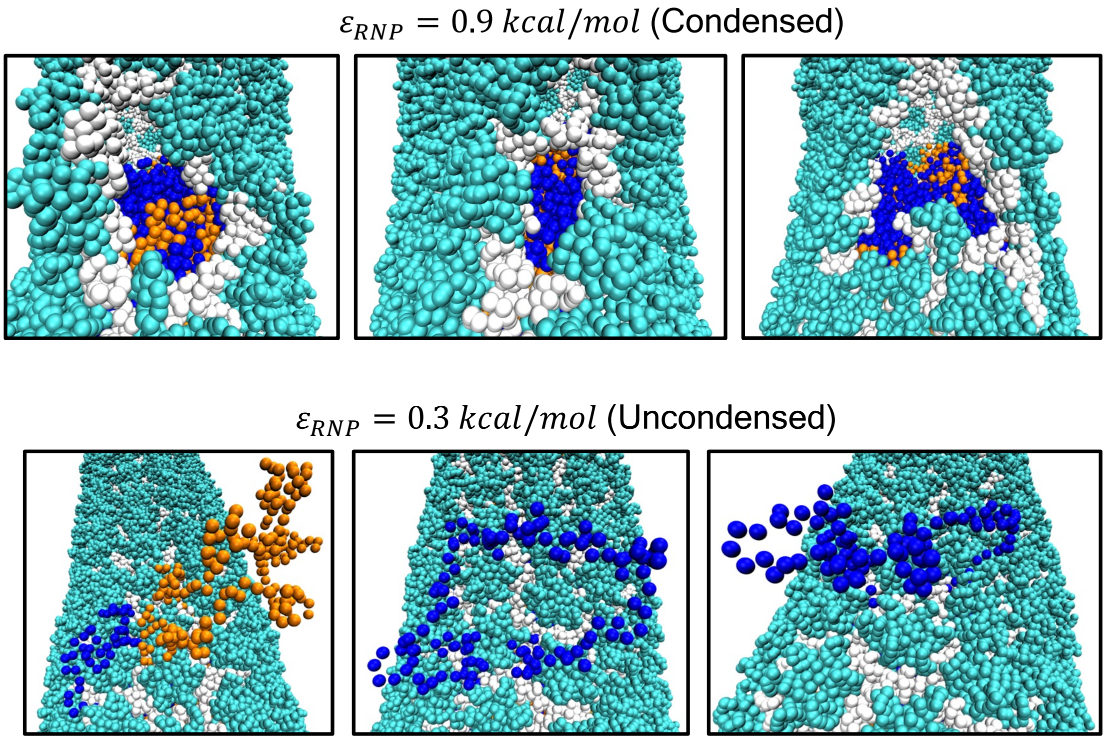


**Fig. S14. Snapshots of the rupture in the capsid lattice and extrusion of the RNP chains for the pill-shaped capsid.** For $\varepsilon_{RNP}$ = 0.9 kcal/mol (upper panel), condensation of the RNP complex creates rupture of the capsid lattice in the vicinity of the globular condensate. For $\varepsilon_{RNP}$ = 0.3 kcal/mol (lower panel), the segments of the uncondensed RNP complex protrudes out of the capsid lattice from the CA lattice rupture.

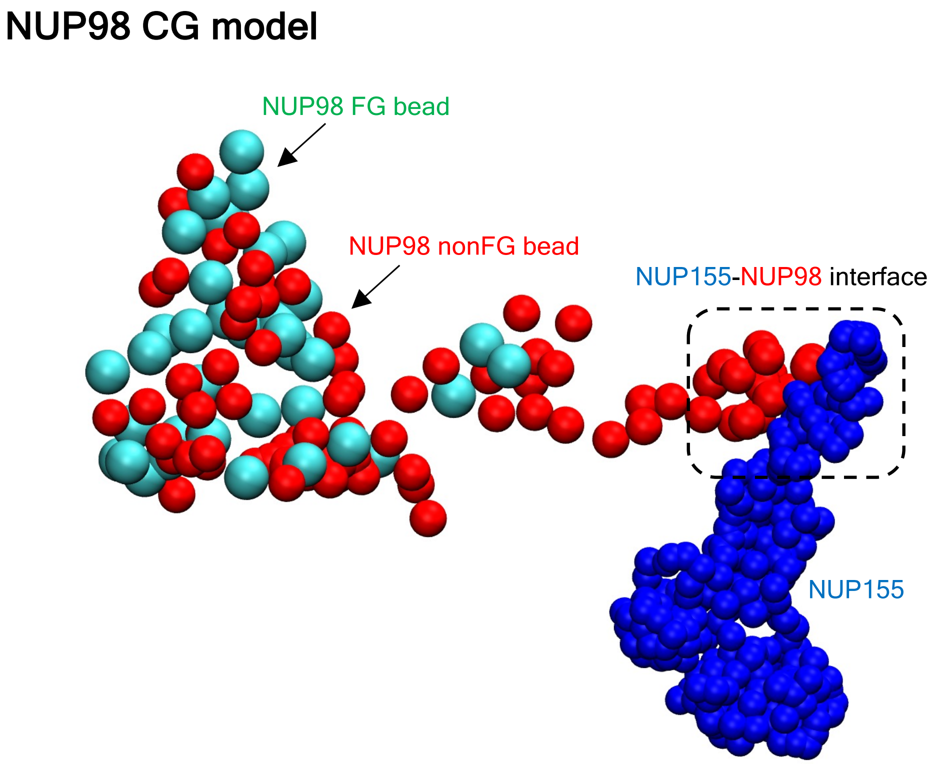


**Fig. S15. CG model of disordered NUP98.** The FG and nonFG CG beads of a NUP98 chain is shown in cyan and red spheres respectively. We also highlight the NU155-NUP98 binding interface based on the PDB: 7R5J (7). NUP155 is shown in blue spheres. The snapshot corresponds to $\varepsilon_{NUP98}$ = 0.3 kcal/mol.
